# Multifractal roots of suprapostural dexterity

**DOI:** 10.1101/2020.07.17.209502

**Authors:** Damian G. Kelty-Stephen, I-Chieh Lee, Nicole S. Carver, Karl M. Newell, Madhur Mangalam

## Abstract

Visually guided postural control emerges in response to task constraints. Task constraints generate physiological fluctuations that foster the exploration of available sensory information at many scales. Temporally correlated fluctuations quantified using fractal and multifractal metrics have been shown to carry perceptual information across the body. The risk of temporally correlated fluctuations is that stable sway appears to depend on a healthy balance of standard deviation (*SD*): too much or too little *SD* entails destabilization of posture. This study presses on the visual guidance of posture by prompting participants to quietly stand and fixate at distances within, less than, and beyond comfortable viewing distance. Manipulations of the visual precision demands associated with fixating nearer and farther than comfortable viewing distance reveals an adaptive relationship between *SD* and temporal correlations in postural fluctuations. Changing the viewing distance of the fixation target shows that increases in temporal correlations and *SD* predict subsequent reductions in each other. These findings indicate that the balance of *SD* within stable bounds may depend on a tendency for temporal correlations to self-correct across time. Notably, these relationships became stronger with greater distance from the most comfortable viewing and reaching distance, suggesting that this self-correcting relationship allows the visual layout to press the postural system into a poise for engaging with objects and events. Incorporating multifractal analysis showed that all effects attributable to monofractal evidence were better attributed to multifractal evidence of nonlinear interactions across scales. These results offer a glimpse of how current nonlinear dynamical models of self-correction may play out in biological goal-oriented behavior. We interpret these findings as part of the growing evidence that multifractal nonlinearity is a modeling strategy that resonates strongly with ecological-psychological approaches to perception and action.

## 1. Introduction

### 1.1. Stability of suprapostural visual activities at longer scales rests on fluctuating behavior at shorter scales

Standing quietly and focusing on a target in front of us is the preamble to various coordinated behaviors —we lean forward and reach or track the target’s progress and bat it away. Less recognized is the fact that this starting position is not merely the preamble to action but is already a rich wellspring of the action itself, exhibiting a continuous stream of intermittent fluctuations. These fluctuations in our bodily center of mass (CoM) and center of pressure (CoP, where ground reaction forces meet our lower extremities) are crucial to maintaining a quiet stance unless they pitch the CoP beyond the base of support (Winter, 1995; Zatsiorsky & Duarte, 1999). These fluctuations offer the body a subtle and flexible command of the mechanical surface underfoot (Chen et al., 2008; Rajachandrakumar et al., 2018), exemplifying long-respected proposals that what looks ‘noisy’ can actually stabilize nonlinear-dynamical systems (Shinbrot and Muzzio, 2001; Turing, 1952). Critically, what looks noisy may instead be richly structured movement variability (Newell and Corcos, 1993; Riccio, 1993; Slifkin and Newell, 1999, 1998; Stergiou et al., 2006)—with richness entailing high-dimensional nonlinear dynamics (Lovejoy, 2019; Schertzer and Lovejoy, 2004), enabling movement systems to sustain their dexterous contact with task context through multifractal foundations (Cavanaugh et al., 2017). This work examines how the visual precision demands reshape the pattern of interactions amongst fractal and multifractal fluctuations in postural control.

We begin with Section 1.1. to define some of the technical terms ahead. We take up these terms beginning in Section 1.2. to develop our rationale motivating this empirical work.

### 1.1. Preliminary definitions of terms for describing movement variability: monofractality, multifractality, and flow of mono- or multifractal fluctuations

At the outset, we wish to define these terms so that any new readers can understand what is at stake in this portrait of movement variability. Briefly, ‘fractal fluctuations’ refer to time series with temporally-correlated sequencing, meaning that each current value is relatively more predictable from previous values than one might find in a completely random sequence (e.g., in additive white Gaussian noise). Additive white Gaussian noise is classically ‘uncorrelated,’ implying no statistical relationship across time between its consecutive values. Specifically, fractal fluctuations entail a temporal correlation in which the variance of current values is strongly correlated with variance of recent values and in which the correlation between the present and past drops off very slowly. Fractal fluctuations present an intriguing case of a non-zero, non-vanishing correlation between current variance and variance in the most distance past. Hence, fractal fluctuations reflect a case of ‘long memory’ in a time series. The clearest signature of this long memory is when the *SD* of a measured time series grows relatively fast. For white noise series without long memory, *SD* grows according to the square root of time (i.e., time with exponent 0.5). For fractally-fluctuating series, *SD* grows linearly with time (time raised to exponent 1). The exponent on time will appear in this work as a ‘Hurst exponent’ *H* abbreviated as ‘ *H*_fGn_,’ because the first step of fractal analysis is to assume that fractal (*f*) fluctuations appear as Gaussian noises (Gn). Hence, monofractal evidence diagnoses ‘fractal Gaussian noise’ (fGn) in terms of values of H_fGn_ deviating away from 0.5 and closer to 1.

Movement variability often exhibits fractal fluctuations in response to monofractal analyses that test for only one fractal exponent *H*_fGn_ at a time. However, with multiple monofractal analyses, movement variability exhibits multiple *H*_fGn_. So obviously, movement variability is not ‘monofractal.’ Using multiple monofractal analyses reveals the texture hidden in the very many degrees of fractal patterning across space and time in the movement system. Hence, movement variability is ‘multifractal,’ which can be analyzed as the temporal or spatial variation of monofractal evidence.

A growing body of evidence suggests that movement variability exhibits a flow of fractal and multifractal fluctuations from one anatomical location or form into another (e.g., Mangalam, Carver, et al., 2020a, 2020b; Stephen et al., 2012). That is, a single case of monofractal and multifractal fluctuations in one part of the body is just the tip of the metaphorical iceberg. This ‘flow’ indicates that we can statistically test how increases in *H*_fGn_ at one location or one point in time predicts subsequent changes in *H*_fGn_ in the same or a different location. We can equally well statistically test how increases in multifractal variety predict subsequent changes in multifractal variety. This work will open up a field of inquiry where we can test for relationships between monofractal estimates (*H*_fGn_), multifractal estimates, and *SD* of postural fluctuations—each of these measures of movement variability can change, and those changes can predict subsequent changes in each other measure of movement variability. The present work will examine how task constraints of quietly standing and visually fixating at a distance prompt different relationships amongst these aspects of postural control.

### 1.2. Stability of suprapostural visual activities at longer scales rests on fluctuating behavior at shorter scales

Eyes lace the postural system into the visual field, locking focus on a distant point in the world; this lock is fluid, nonetheless. Fixations are regularly recognized as a class of movement. Just as upright posture is never entirely still, the ‘fixations’ of the eyes are not stasis (Montesano et al., 2018). Fixations consist of a deep, fluctuating foundation of smaller movements called ‘microsaccades’ (Meyberg et al., 2017). These microsaccades stabilize images that would otherwise fade on a static retina (Martinez-Conde et al., 2013), thus playing a similar role to the visual system that fluctuations play for the postural system. Indeed, the exploratory role of fluctuating movements extends to the bodily extremities besides and intermediating between upright posture and vision (Murnaghan et al., 2016), suggesting a vital role for postural sway in supporting visual perception (Stoffregen, 2010; Stoffregen et al., 2000, 1999). Hence, even when quietly standing and fixating at a distant target, the body is coursing with fluctuations ferrying information across the body.

Visually engaging with the world contributes to the fray of ongoing bodywide work to stay on task and dexterously responsive to context. Fixations require precision like all other movements, stabilizing our position in the world. By ‘visual precision demands’ we do not mean the laboring of visual-cognitive mechanisms (Wang et al., 2013), but instead the body-wide muscular work maintaining fixations straight ahead at a static stimulus. This term can be undressed of possible confusion with the visual-cognitive workload by substitution with ‘ocular precision demands,’ which does not necessarily denote maintaining fixation to single body part. Indeed, no less than the postural system seeking stability amidst wide-ranging constraints, the precision needed to maintain fixation is the work of the entire body positioned amongst task constraints like ambient light and surface (Cachinero-Torre et al., 2017; Kaldy and Blaser, 2020).

Postural fluctuations produce translations and rotations of visible surfaces specific to the objects’ spatial relationships in the visual field. These movement-induced translations and rotations compose an optic flow that provides visual support for the subsequent movement. Thus, subtle sway offers the sighted organism a rich source of information about objects’ layout in the visual field. The visual layout itself then presses the postural system into a poise for engaging with objects and events. Visual targets appearing closer to or farther from the looking postural system impose retina-specific oculomotor convergence constraints (Jaschinski-Kruza, 1991; Jaschinski, 2002) and chromatic aberration (Párraga et al., 2002). Consequently, changes in a target’s size relative to other aspects of the visual field entail optic flow (Fig. 1). These multiscale factors affect postural stability, reflecting changes in postural configurations and the flow of information needed to organize these postures. Although how these multiscale factors enact their effects remain controversial, observations of across-scale interactions in sway align with the proposal that the coordination of posture is a fluid, cascading process in which relatively coarse, longer-scale constraints (e.g., the task) redirect progressively finer-grained motoric degrees of freedom without stipulating them (Latash, 2020, 2012). This cascade operates through the loose nesting of finer motoric details within larger motoric degrees of freedom (e.g., motor neurons within bundles of muscle fiber, muscles within joint-centered synergies, and so on) that allow dexterous behavior to adapt flexibly to constraints changing on the way to task completion.

**Figure 1.**
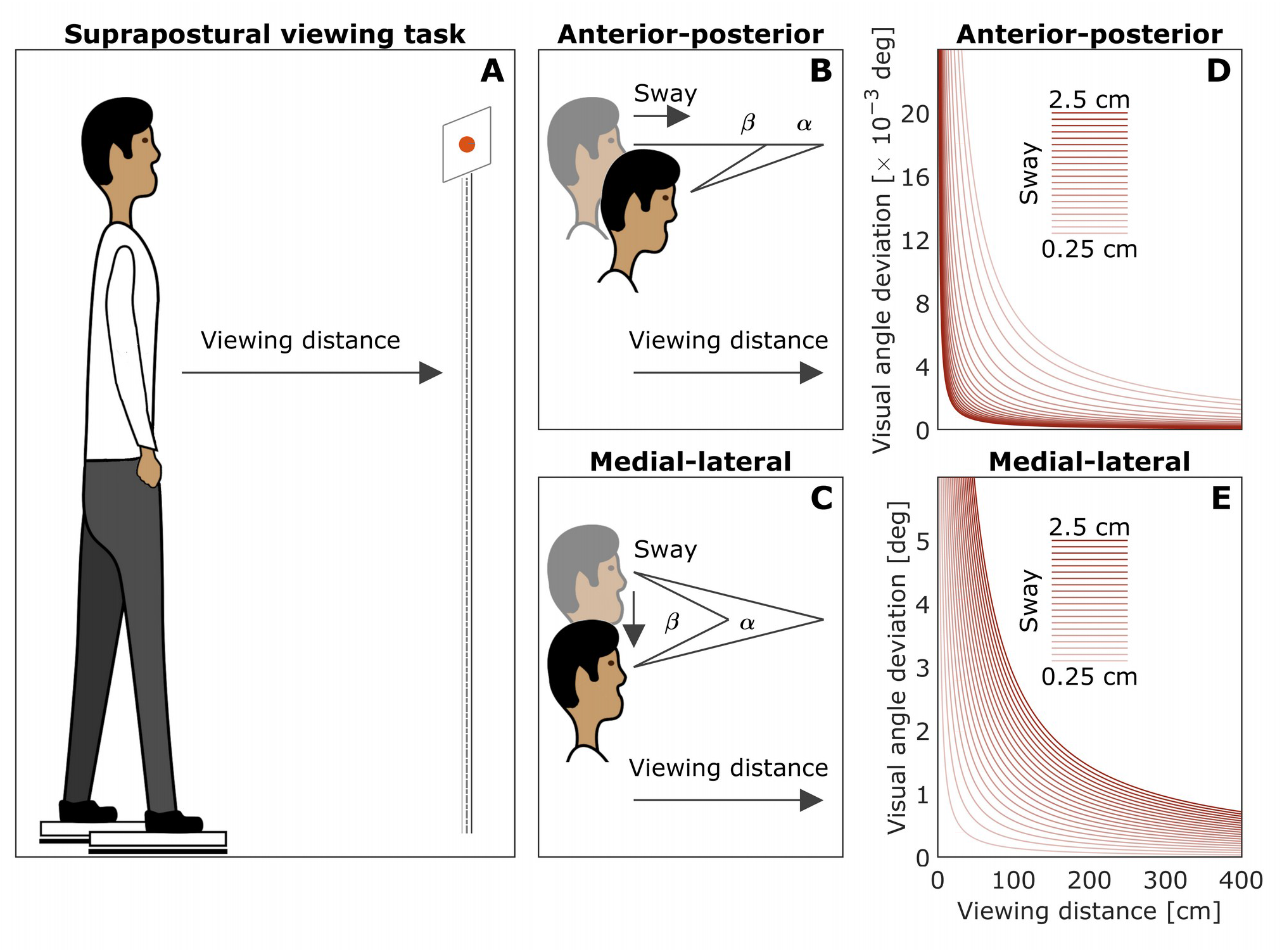
Schematic of the task and effects of eye-to-target distance on postural sway. (**A**) The suprapostural viewing task of standing quietly with the eyes fixated at a distant visual element. (**B**, **C**) Visual angle gain for short vs. long eye-to-target distances along the anterior-posterior (*AP*) and medial-lateral (*ML*) axes. (**D, E**) Visual angle gain as a function of eye-to-target distance for different sway magnitudes. Closer targets increase *AP* sway, and farther targets increase *ML* sway.

### 1.3. Visual precision demands on postural stability could reshape intrapostural interactivity

Movement depends on the bodywide network of connective and nervous tissues, forming flexible relationships that balance tensions with compressions at multiple scales (Cabe, 2018; Ingber, 2006, 2008, 2010; Nelson et al., 2005; Schleip et al., 2014; Turvey & Fonseca, 2014). This organization entails a multifractal geometry that embodies multiple scale-invariant patterns of behavior (e.g., microsaccades within the saccades that intersperse larger saccades by the eye and turns by the head) across time and space. CoP multifractality has repeatedly borne a consistent relationship to perceptual judgments of visual and haptic stimuli (Doyon et al., 2019; Hajnal et al., 2018; Mangalam et al., 2020c; Mangalam and Kelty-Stephen, 2020; Palatinus et al., 2014, 2013). Research on perceptual tasks (e.g., manually wielding an object to judge its heaviness or length has shown that a bodywide flow of multifractal fluctuations precedes and shapes the verbal articulation of perceptual judgments (Mangalam et al., 2020a, 2020b). Hence, the flow of multifractality within posture supports information flow and might provide a glimpse of the control policy emerging from the bodily situation in task constraints.

Indeed, the intermodal perception supporting standing and fixating makes an inquiry into multifractal roots particularly valuable. Multifractal geometry appears not simply important for wholesale pickup of information but is specifically predictive of how different tissues of the body coordinate different aspects of information about the task context (Carver et al., 2017; Doyon et al., 2019; Hajnal et al., 2018; Kelty-Stephen and Dixon, 2014). Multifractal aspects of sway also affect the coordination of mechanical and visual information as people stand still while visually fixating. Changing the movement system’s relationship to visual information (e.g., increasing precision demands or closing eyes) while standing on the same mechanical surface could prompt the body to arrive at new body-wide coordination. We expect that changing this coordination of visual and mechanical information would depend on multifractal foundations. Hence, this work investigates fractal evidence that would speak to ongoing deliberations about clinical applications to supporting upright stance (e.g., Hove et al., 2012; Rodger & Craig, 2016; Uchitomi et al., 2013). It also connects fractal results and discourse to multifractal processes underlying fractal evidence and supporting the more generic intermodal coordination.

### 1.4. Visual precision demands for fixating across different viewing distances could reveal patterns of intrapostural interactivity

#### 1.4.1. The task

This work is a reanalysis of a previous study that manipulated viewing distance and measured CoM and CoP (Lee et al., 2019). Healthy adults stood and maintained a quiet stance for 120 s under six different conditions: a control condition with eyes closed and five conditions with eyes open and fixating on a red laser point projected on surfaces at 20, 50, 135, 220, and 305 cm distance. These five non-equally progressively spaced distances captured the power-law like relationship between viewing distance and resulting angular deviation under the assumption of 2-cm postural sway in either *AP* or *ML* direction (Figs. 1D and 1E). The authors referred to this nonlinear relationship between viewing distance as ‘visual angle gain’ and used it to express the range of optic flow expectable from standing as still as possible. The task follows in a long, productive tradition of examining how visual perception of near or far targets influences posture (Bles et al., 1980; Lee & Lishman, 1975; Mayo et al., 2010; Munafo, Curry, et al., 2016; Munafo, Wade, et al., 2016; Stoffregen et al., 1999, 2000).

The original data collection allowed Lee et al. (2019) to examine the frequency of intermittent postural corrections observed through increased bursts of posture–head movement coordination. Critically for our reanalysis, this task and dataset offer the opportunity to examine the novel issue of intrapostural interactivity. Indeed, postural sway can produce unintended movements that nonetheless serve the perceptual goals, and as Lee et al. (2019) themselves demonstrated, the task-oriented aspect of these movements may manifest in any of many different statistical measures of postural sway (Munafo et al., 2016b, 2016a; Palatinus et al., 2014, 2013). We can undoubtedly examine concurrent correlations amongst these different statistical measures, but in this work, our curiosity is specifically about the exchange of fluctuations as they change with the visual precision demands. To our knowledge, the long tradition of studying postural sway in visual tasks does not yet include causal modeling of movement variability to determine which fluctuations precede and potentially influence subsequent changes in other fluctuations.

For our purpose, the two explicit precision demands from Lee et al.’s (2019) allowed an elegant experimental clarity. Asking participants to fixate their gaze and minimize postural sway nearly froze not one but two linkages to the task context, leaving the participant’s movement system taut between static visual and static mechanical supports. This manipulation reflects the ecological fact that sway is never due to movement in the visual or mechanical stimulation, and changes in postural sway are most clearly due to the ongoing cooperations and competitions amongst postural degrees of freedom. Hence, the double-precision command offers a unique opportunity to delve into intrapostural interactivity. The manipulation of viewing distance is effectively a manipulation of the visual precision demands. The 50-cm distance is within an ideal range to focus on red light (Párraga et al., 2002) and prompts the least straining oculomotor convergence (Jaschinski-Kruza, 1991; Jaschinski, 2002). We predicted that increasing precision demand by requiring viewing distances farther from 50-cm viewing distance would accentuate relationships between sway *SD* and fractality (i.e., *SD* and fractality of postural fluctuations).

#### 1.4.2. Modeling strategy

To accomplish our goal, we reanalyzed data from Lee et al. (2019b) in three steps: (i) carving individual trials under each viewing condition into non-overlapping sub-trial segments, (ii) estimating *SD* and fractal temporal structure within each segment, and (iii) using vector autoregression to test the interactions among these descriptors across segments within each trial.

*SD* has been an intuitive way to model stability, but as noted above, it has only limited heuristic value as an inverse measure of stability. Fractal and multifractal indices have drawn attention in postural-control research only as a point of contact with nonlinear dynamics, cascades, and self-organized criticality. The fractal temporal structure of bodywide fluctuations has been implicated in predicting variability in perceptual judgments or responses capable of deviating from instructed or intended accuracy/veridicality. Whatever value *SD* serves for postural control goals, we can take *SD* as an analogous measure of perceptual outcome: sway *SD* serves as a measure of how well participants were able to follow the instructions to minimize their sway. Minimizing sway is by no means the same as an explicit perceptual judgment, but like a perceptual response to instructions, it can be done with varying degrees of accuracy.

Our choice of descriptors is not entirely divorced from that of Lee et al.’s (2019) choice of modeling both *SD* and correlation dimension. The correlation dimension is a fractal dimension, but instead of emphasizing temporal sequence, it emphasizes spatial recurrence of a phase-space trajectory. For instance, the power-law scaling of %recurrence with a radius in recurrence quantification analysis estimates the correlation dimension. Thus, our choice of dependent variables is not entirely novel, but it differs to align this work with other postural and non-postural cases in which fractal fluctuations support perceptual responses.

##### 1.4.2.1. Why we are modeling multifractality as temporal variation in fractal analysis, instead of estimating only a multifractal spectrum

Our choice to model the relationship between *SD* and fractality sits within our foregoing interest in applying the multifractal formalism to perception and action. In the context of discussing multifractality, the term ‘fractal’ does entail ‘monofractality,’ and the prefix ‘mono-’ does mean ‘not multi-.’ The use of monofractality after just having trumpeted multifractality may seem jarring to seasoned readers. However, our choice here has three points of motivations.

First, to our knowledge, all current empirically supported claims implicating multifractality as an important factor in perception and action began with or were inspired by monofractal analysis as a preliminary step (Mangalam et al., 2020c, 2020a; Palatinus et al., 2013). So, because we ask a new question about postural control while fixating at a distant target and not merely aiming to examine multiple dependent measures about postural sway separately, we hew to the tradition of modeling the predictive effects of variation in monofractality. It is worth noting that, given the long shadow cast by information-processing accounts offering more familiar, more intuitive models of goal-directed behavior, many readers and stakeholders in this literature are still coming to grips with monofractality. Leaping straight to multifractal analysis is a step of complexity that might well bury any usefulness of this work.

‘Stochastic resonance’ is an essential example of monofractal roots of postural control. The clinical application of ‘stochastic resonance’ technologies aim to enhance movement dexterity using mechanical or acoustic stimulations patterned according to white-noise or 1/*f* noise. Both are defined as the lack or presence of monofractality. We are excited to see that research into this application of temporally-structured stimulation has recently begun to look beyond lack-vs.-presence of monofractality and, in the vein of recent theorizing (e.g., Cavanaugh et al., 2017; Stergiou et al., 2006), to recognize that the fluctuations supporting the movement are not ‘noise’ at all (Hove et al., 2012; Rodger and Craig, 2016; Uchitomi et al., 2013). Indeed, our modeling strategy resonates with these calls, suggesting that we may need to give voice to postural variations in monofractal estimates rather than a simple lack-vs.-presence dichotomy. Nonetheless, the fact remains that monofractality remains foundational to research interest in postural fluctuations. Interest in multifractality strikes us as the need for the judicious situation of monofractal analysis sooner than a reason to eschew monofractal analysis.

Second, variation in monofractality is itself multifractality. Temporal or trial-by-trial variation of monofractal evidence is one of the most accessible ways in which to observe and communicate multifractality (Ihlen, 2012). Multifractality can be estimated as a spectrum of fractal-scaling dimensions, and preceding research on postural control has examined multifractality through this very multifractal spectrum (Koslucher et al., 2016; Munafo et al., 2016a). However, despite this constructive precedent, we still find value in examining multifractality as the variations of monofractal evidence for a third reason.

Finally, although the multifractal spectrum results from variations in monofractal evidence across time, it is not informative about the exact sequence of those variations. Variations of monofractality deliver two important insights that a single multifractal spectrum aggregating those variations could not. The first is a direct view of the multifractal spectrum width, albeit spread out over time. The second is a direct view of how the increases and reductions of monofractality could predict subsequent changes in *SD*. Similar-sized changes in monofractality in opposite directions would give us the same multifractal spectrum. The direction of change in monofractality could be predictive of subsequent *SD* that multifractal spectrum width itself could not predictive. If that were the case, then leaping to multifractal spectrum at the cost of examining variation in monofractal evidence would risk a false-negative result.

##### 1.4.2.2. Why we are modeling multifractal-spectrum width as well to test whether nonlinear interactions across scales support the subsequent changes in SD and monofractal variation

Far from suggesting that estimating the multifractal spectrum is by itself a poor choice, this work uses it to quantify the strength of nonlinear interactions across scales. The strength of nonlinear interactions across scales can be quantified by comparing a measured series’ multifractal spectrum width to the multifractal spectrum widths of surrogate series, preserving the linear structure of the original measurement but destroying the originally measured sequence. Indeed, the *t*-statistic comparing original multifractality to surrogate multifractality (henceforth, ‘multifractal nonlinearity’ *t*_MF_) appears to be negatively correlated with concurrent *SD* in precision tasks and in postural sway (Bell et al., 2019; Jacobson et al., 2020; Kelty-Stephen, 2018).

Although we can estimate monofractality and multifractality separately, the control of goal-directed behavior reflects a balance between the two. The multifractal spectrum width reflects the flexible capacity for the body to embody multiple scale-invariant patterning, but this width itself only speaks to variety and not to exact direction of relationships across time. We expect bidirectional relationships between monofractal evidence and multifractal nonlinearity. (Note: we refer here to ‘monofractal evidence’ to reflect the fact that only conducting monofractal analyses can generate monofractal evidence without any indication that the analyzed system is itself monofractal and not multifractal). Changes in monofractal evidence do entail changes in multifractal structure (Ihlen, 2012), and thus changes in that monofractal evidence should predict subsequent changes in ongoing multifractal nonlinearity. Multifractality gains its causal force in shaping goal-directed behaviors not simply through the diversity of monofractality but also from the fact that the individual monofractality specify a degree of correlation. Monofractal temporal correlations have a valence to them, with greater or weaker correlations indicating the longer or shorter propagation of events across time scales (Kuznetsov and Wallot, 2011). Indeed, we might conceive multifractality as a reservoir of flexible possibility for nonlinear interactions across timescales, but temporal correlations are an important, supportive constraint in guiding multifractal behavior (Cavanaugh et al., 2017; Wallot et al., 2015).

At the outset (Section 1.2.), we noted that multifractal fluctuations bore a specific relevance to predicting how movement systems coordinate intermodal perception as in standing and visually fixating. We expect that multifractal nonlinearity is the foundation supporting monofractal variation and supporting any relationship between monofractal variation and *SD* of sway. Hence, we will aim in this study to, first, establish that monofractality might hover around a set point as posture ranges between the center and the margin of the support surface and, second, that increases in monofractality might precede subsequent reductions in *SD*. However, the concluding goal of this work is to demonstrate that multifractal nonlinearity is a crucial factor in both these features of monofractality in postural control.

To clarify, and to foreshadow our hypotheses, we expect that multifractal nonlinearity is the crucial guarantee of the capacity for monofractal variation. And we further expect that any effects of monofractality on *SD* of postural sway depend on nonlinear interactions across scales. That is, despite their being conceptually separate, multifractal nonlinearity and monofractality hang together statistically. So, we expect that multifractal nonlinearity will predict subsequent reductions in *SD*. However, we also predict that any evidence of monofractality predicting subsequent reductions in *SD* may be better explained by multifractal nonlinearity.

##### 1.4.2.3. What multifractal nonlinearity could mean for intrapostural interactivity supporting the suprapostural task of quietly standing under various visual precision demands

Although variation in monofractal evidence indicates interactions across scales that support dexterity, multifractal nonlinearity is a stronger operationalization of these interactions (Kelty-Stephen and Wallot, 2017). Hence, we propose that multifractal nonlinearity is more central than monofractal evidence to the task- or context-dependence of goal-directed actions. This task-dependence appears here as visual precision demands. We introduced above the idea that visual precision demands would moderate the association between prior monofractal evidence and subsequent reduction in *SD* of postural fluctuations. Certainly, in the absence of estimates of multifractal nonlinearity, monofractal evidence will be a strong first pass at modeling organismal sensitivity to task constraints. However, when we introduce multifractal nonlinearity into the model, we expect this sensitivity to task constraints much more in multifractal nonlinearity than in monofractal evidence.

We expected that incorporating multifractal nonlinearity into the vector-autoregressive relationships would radically reshape our view of the intrapostural interactivity supporting postural control. First, because multifractality is itself the generalization of monofractal structure, we expected it to show more refined sensitivity to task constraints. For instance, whereas we predicted that the effects of monofractal evidence on subsequent *SD* would be sensitive to visual precision demands, we expected that multifractal nonlinearity would better predict these effects of visual precision demand and, further, predict effects of straightforward availability of visual information (i.e., eyes-open vs. eyes-closed) on subsequent *SD*.

Multifractality increases as the variety of monofractal patterns in movement exceeds the monofractal variety in linear models of that movement. Hence, multifractal nonlinearity is an asset in tasks that require ongoing exploration (Booth et al., 2018; Carver et al., 2017) but a liability in tasks that require freezing out degrees of freedom (Bell et al., 2019; Jacobson et al., 2020; Kelty-Stephen, 2018). We expected multifractal nonlinearity to show both these faces in this work: as a stabilizing factor with increasing visual precision demands and destabilizing factor under abrupt changes in the informational support for quiet stance. Specifically, visual precision demands should accentuate an *SD*-reducing effect of multifractal nonlinearity—increasing the visual precision demands might prompt a reduction of variability in average positions or trajectories, but in order to rein in the SD of postural positions (i.e., of the center of pressure, CoP), the movement system still needs exploratory behaviors to use the precision demand. Indeed, precision-related reductions in *SD* of position can coincide with stronger signatures of complexity, suggesting that exploratory behavior does not live solely in the bulk of movement variability but also in its patterning (Haddad et al., 2010, 2008). After all, extrinsic constraints from the task context are rarely stipulated impositions and much more often available supports for organisms to engage with through exploration or not engage at all (Stoffregen et al., 2017). Precision demands can be no different but are more likely available features of the context with effects only contingent on the organism’s exploratory behaviors. We expect that organisms can meet progressively stronger precision demands only through progressive—and prior—increases in their exploratory behaviors’ multifractal nonlinearity.

Closing the eyes should introduce an *SD*-increasing effect of multifractal nonlinearity. Closing eyes abruptly changes the visual support for posture (Stoffregen, 1986), destabilizing the movement system, and the priority for the postural system to maintain quiet stance is sooner to damp out fluctuations and freeze degrees of freedom. Hence, if we destabilize posture, postural sway (e.g., as measured by *SD* of CoP fluctuations) generally increases with greater multifractality (Kelty-Stephen, 2018). These prior observations include a failure to find a concurrent correlation of multifractal-spectrum width with *SD* of CoP position. Multifractal variability has elsewhere varied inversely with stable postural outcomes, with narrower multifractal spectra appearing in younger adults and in participants less likely to experience motion sickness (Koslucher et al., 2016; Munafo et al., 2016a). However, previous work has never before integrated the *t*_MF_ statistic, vector-autoregressive effects of one variable on subsequent values of another, and the potential moderation by visual precision. Hence, the present test of the effects of multifractal nonlinearity on subsequent *SD* of CoP fluctuations with greater precision demands remains so far untested in the literature.

As the second entailment of including multifractal nonlinearity is the possibility of predicting the apparently ‘self-correcting’ aspect of monofractal evidence. For example, as noted above, the need for postural sway to reverse course from the base of support’s margins has been a long-standing explanation of why postural control shows a transition from stronger and weaker monofractal evidence by turn (Collins and De Luca, 1995, 1993). That is, we expect that increases in multifractal nonlinearity would prompt oscillations in subsequent monofractal evidence. Changes in monofractal evidence may well contribute to multifractal nonlinearity, but because of the centrality of multifractal nonlinearity to task-sensitivity, we suspect that the effects of monofractal evidence on subsequent multifractal nonlinearity would be strongest in less demanding tasks. The corollary is that greater visual precision demands would minimize the contribution of monofractal evidence to multifractal nonlinearity, emphasizing multifractal nonlinearity to the exclusion of any effects of monofractal evidence.

The final entailment of including multifractal nonlinearity is a possible displacement of the effects linking monofractal evidence to subsequent *SD* and linking *SD* to subsequent monofractal evidence. Previous work does not predict whether the effects of monofractal evidence on subsequent *SD* might go to zero or even reverse. However, given the more central and substantive meaning of multifractal nonlinearity supporting dexterous goal-directed behavior, we expected multifractal nonlinearity to influence subsequent *SD* and, furthermore, show subsequent changes due to prior changes in *SD*.

In summary, elaborating from mono-to multifractal formalisms will allow us to delve deeper into the multifractal structure supporting postural control. This move aims to situate monofractal evidence—which can be modeled entirely linearly—within a broader framework of nonlinear relationships. Instead of simply reporting a correlation table of concurrent relationships, we aim to model how SD, monofractal evidence, and multifractal nonlinearity predict subsequent changes in one another.

#### 1.4.3. Hypotheses

##### Hypothesis-1: Weaker intrapostural interactivity while quietly standing with eyes closed

First, respecting the evidence that maintaining fixation recruits visual precision demands that can perturb posture (Fiorelli et al., 2017; Lê and Kapoula, 2008), we expected that the addition of a fixation task to quiet standing would generally accentuate intrapostural interactivity, suggesting that quietly standing with eyes closed would show weaker evidence of intrapostural interactivity.

##### Hypothesis-2: Self-correction of sway monofractality across 10-s segments

Postural sway is more correlated at shorter timescales on the order of 10 s and more anticorrelated over longer timescales (Collins and De Luca, 1993). For instance, posture roams freely about a fixed average CoP position but self-corrects at the margins of the base of support to maintain a quiet stance (Collins and De Luca, 1995, 1993; Gilfriche et al., 2018). In this sense, we expected that monofractal evidence itself would show similar self-corrections over time, with prior increases in fractal evidence followed by subsequent alternating reductions and increases in monofractal evidence. This self-correcting feature would be a crucial part of any control system in which temporal correlations stabilize sway.

##### Hypothesis-3: Inverse relationship between sway monofractality and SD across 10-s segments

Because monofractality entails long-range temporally organized responsivity to mechanical perturbations (Duarte and Sternad, 2008; Duarte and Zatsiorsky, 2001), we expected that either or both of CoP and CoM monofractality would show an inverse relationship with CoP *SD*. That is, increases in CoP and CoM monofractality would predict subsequent reductions in CoP *SD* (Hypothesis-3a), as well as increases in CoP *SD* would predict subsequent reductions in CoP and CoM monofractality (Hypothesis-3b). This feature could reflect a second key component of task-dependent emergent postural control.

##### Hypothesis-4: The inverse relationship between sway monofractality and SD is stronger with eyes open than eyes closed and becomes progressively stronger at viewing distances closer to 50 cm

The 50-cm viewing distance is within the comfortable viewing distance, ideal for the human eyes’ focus of red light (Párraga et al., 2002), as well as ideal for requiring the least straining oculomotor convergence (Jaschinski-Kruza, 1991; Jaschinski, 2002). Thus, we expected that the 50-cm viewing condition would yield intrapostural interactivity most closely resembling that in the eyes-closed condition. Because 50-cm imposes the weakest precision demand, the effects among CoM monofractality, CoP fractality, and CoP *SD* would show the least differences from the eye-closed condition. We also expected that the increasing visual precision demand as viewing distance deviates from 50 cm would accentuate the inverse relationship between prior monofractality and subsequent CoP *SD*. That is, increases in CoP monofractality would lead to less subsequent CoP *SD* at viewing distances 20 and 135 cm than at viewing distances of 220 and 305 cm.

##### Hypothesis-5: The influence of sway multifractal nonlinearity t_MF_ on subsequent increases in sway SD through the bidirectional relationship of t_MF_ with monofractal evidence H_fGn_

We expected that sway multifractal nonlinearity *t*_MF_ would displace monofractal evidence *H*_fGn_ as the stronger predictor of subsequent reductions in *SD* with greater precision demands. Specifically, we expected that increases in *t*_MF_ would predict later reductions in *SD* and progressively larger reductions with progressively stronger visual-precision demands (Hypothesis-5a).

We expected that multifractal nonlinearity would also predict a difference between eyes-closed condition and 50-cm viewing condition that multifractality would not. Specifically, we expected that, for the eyes-closed condition, increases in multifractal nonlinearity would predict subsequent increases in *SD* of postural fluctuations (Hypothesis-5b).

We also expected that (i) multifractal nonlinearity would infiltrate the earlier-predicted bidirectional effects only between monofractal evidence and *SD* (e.g., in Hypotheses-3 and −4); (ii) multifractality nonlinearity would predict subsequent oscillatory reversals in monofractal evidence (Hypothesis-5c); (iii) monofractal evidence would predict subsequent increases in multifractal nonlinearity with progressively lower visual precision demands (Hypothesis-5d); (iv) including multifractal nonlinearity in our modeling would reverse the monofractal-only Hypothesis-4 as follows: with the inclusion of multifractal nonlinearity, monofractal evidence would fail to predict subsequent precision-dependent reductions in *SD*, yielding instead potentially only partially negative or potentially positive subsequent changes in *SD* (Hypothesis-5e). Finally, we expected that including multifractal nonlinearity in our modeling would reverse the monofractal-only Hypothesis-3 as follows: with the inclusion of multifractal nonlinearity, prior increases in *SD* would predict subsequent reductions in multifractal nonlinearity—and no longer any subsequent changes in monofractal evidence (Hypothesis-5f).

## 2. Materials and methods

### 2.1. Participants

Seven adult men and nine adult women (*M±1SD* age = 23.8±3.9 years) without any skeletal or neuromuscular disorder voluntarily participated after providing verbal and written consent approved by the Institutional Review Board (IRB) at the University of Georgia (Athens, GA).

### 2.2. Experimental task and procedure

Each participant stood barefoot with one foot on each of two force plates (AMTI Inc., Watertown, MA), 25 cm apart (figure 1*a*). From behind the participant, a laser pen projected a static point-light on the center of a 5×5” white tripod-mounted screen in front of a white visual-field-filling background. The two force plates measured 3D moments and ground reaction forces. The full-body motion of each participant was measured using VICON Plug-in Gait full-body 39 marker set and an 8-camera VICON motion tracking system (VICON Inc., Los Angeles, CA). The kinetic and kinematic data were synchronously sampled at 100 Hz.

Each participant was instructed to maintain a standing position and minimize postural sway while looking at a point projected in front of them at their eye level with their head in a neutral position. Each participant was tested under six different viewing conditions: eyes closed and the point-light point positioned at 25, 50, 135, 220, and 305 cm distance in front. Each participant completed 18 trials (6 conditions × 3 120-s trials) in a single 90-min session with randomized trial order and with breaks on request and between every six trials.

### 2.3. Data processing

All data processing was performed in MATLAB 2019b (Matlab Inc., Natick, MA). The position of the bodily center of mass (CoM) was estimated by submitting segment lengths of the head, trunk, pelvis, and left and right hand, forearm, upper arm, thigh, shank and foot to the equations provided by Zatsiorsky and Seluyanov (Zatsiorsky and Seluyanov, 1985), which yielded a 3D center of pressure (CoP) series describing CoM position along the participant’s anterior-posterior (*AP*), medial-lateral (*ML*), and superior-inferior axes. 3D moments and ground reaction forces measured on each trial yielded a 2D center of pressure (CoP) series describing CoP position along the participant’s AP and ML axes. Over 120 s duration, each trial yielded a 3D CoM series and a 2D CoP series each of 120 s or 12000 samples, divided into 12 segments of 10 s or 1000 samples each. Each segment yielded two 999-sample one-dimensional series: a CoM spatial Euclidean displacement series describing the amplitude of CoM displacement and a CoP planar Euclidean displacement series describing the amplitude of CoP displacement (Figs. 2A to 2F). We estimated the SD of CoP planar Euclidean displacement series (henceforth, CoP-*SD*).

**Figure 2.**
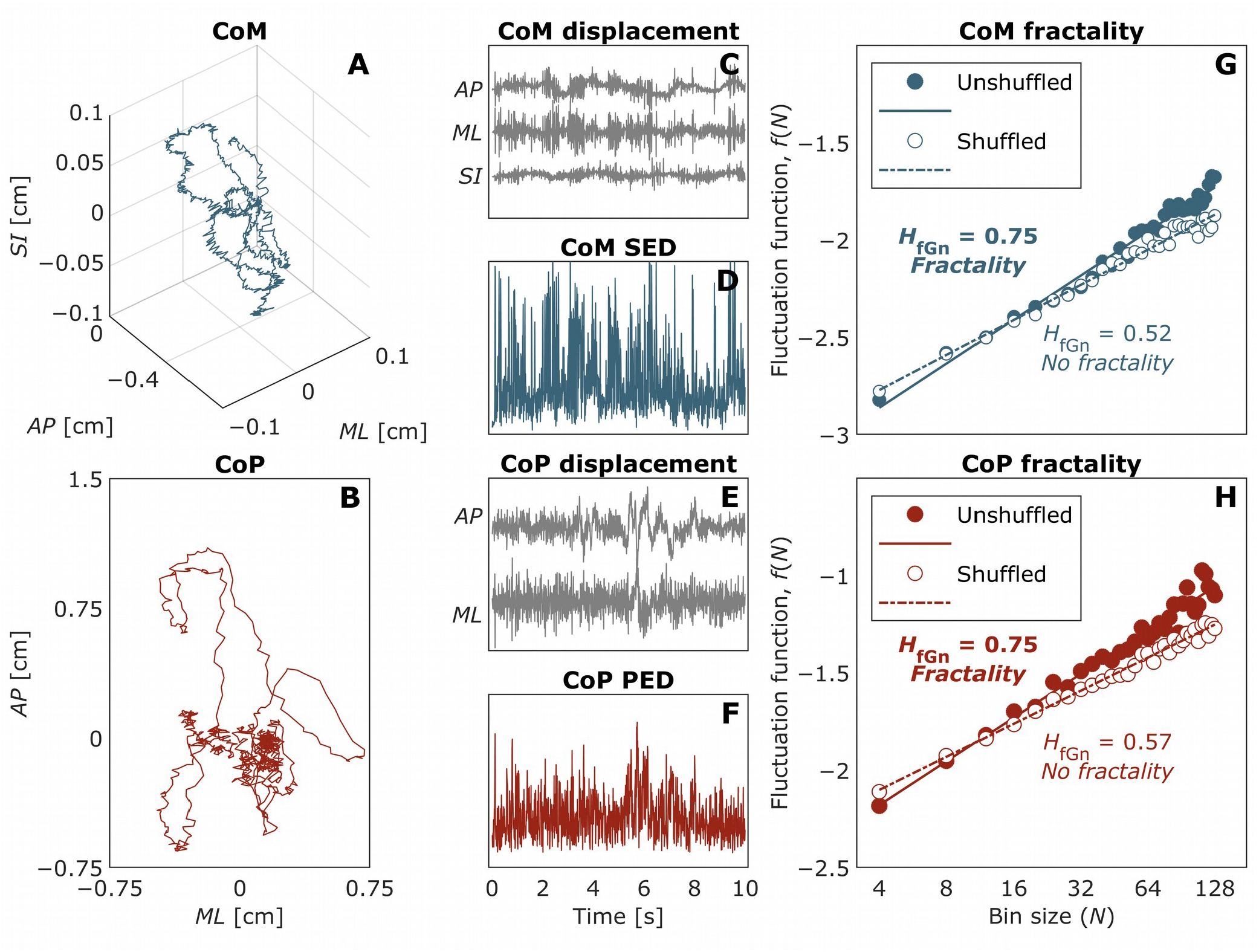
An overview of detrended fluctuation analysis (DFA). (**A, B**) CoM and CoP in a representative 10-s segment. (**C** to **F**) CoM displacement along the medial-lateral (*ML*), anterior-posterior (*AP*), and superiorinferior (*SI*) axes; CoM spatial Euclidean series; CoP displacement along the *ML, AP,* and *SI* axes; CoP planar Euclidean series. (**G, H**) Log-log plots of fluctuation function, f(*N*), vs. bin size (*N*), reflecting the fractal evidence, *H*_fGn_, yielded by DFA. Solid circles and solid trend lines represent *f*(*N*) for the original (unshuffled) series; open circles and dashed trend lines represent *f*(*N*) for a shuffled version of the original series.

### 2.4. Detrended fluctuation analysis (DFA)

DFA estimates Hurst exponent, *H*, describing the growth of root mean square (RMS) fluctuations with time for first-order displacements known as fractional (‘fractal’ for short) Gaussian noises (fGn) (Peng et al., 1995, 1994). First, it integrates time series *x* (*t*) with *N* samples to produce *y* (*t*):

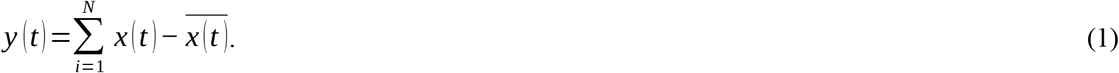

Next, DFA computes RMS residuals from the linear trend *y_n_*(*t*) over nonoverlapping *n*-length bins of *y*(*t*) to build a fluctuation function *f*(*N*):

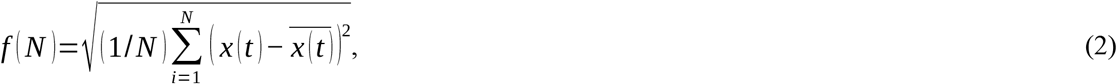

for *n<N*/4. On standard scales, *f* (*N*) is a power law, that is, 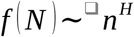, where *H* is the scaling exponent. *H* is estimated as the slope of *f* (*N*) in log-log plots:

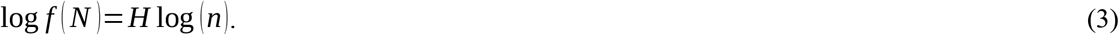

DFA estimated *H_fGn_* for the original version (i.e., unshuffled) and a shuffled version (i.e., a version with the temporal information destroyed) of each segment of CoM spatial-displacement series (henceforth, CoM-*H*_fGn_) and CoP planar-displacement series (henceforth, CoP-*H*_fGn_) over the following bin sizes: 4, 8, 12,… 128).

*H*_fGn_ for all original series fell in the fractal range (0.5 < *H*_fGn_ < 1; Fig. 2G and 2H) and significantly exceeded *H*_fGn_ for all corresponding shuffled series (*P*s < 0.0001), indicating evidence of monofractality in both CoM and CoP (e.g., Figs. 2G and 2H). Crossovers between shorter- (bin sizes: 4, 8, 12,… 64) and longer-scale (bin sizes: 4, 8, 12,… 128) behavior exhibited no reliable differences, as *H*_fGn_ estimates from only shorter-scale behavior correlated strongly with *H*_fGn_ estimates from the entire DFA fluctuation function (CoM: Spearman’s *ρ*s = 0.96, 0.91, 0.94, 0.94, 0.93, and 0.94 for the eyes-closed and 25-, 50-, 135-, 220-, and 305-cm viewing conditions, respectively, *P*s < 0.0001; CoP: *ρ*s = 0.96, 0.89, 0.93, 0.92, 0.91, and 0.94, *P*s < 0.0001).

### 2.5. Direct estimation of multifractal spectra

The multifractal spectrum is a relationship between two fractal dimensions *α* and *f* (*α*). One multifractal elaboration of DFA uses DFA-estimated *H*_fGn_ to estimate both *α* and *f* (*α*) (Kantelhardt et al., 2002). However, we use Chhabra and Jensen’s (1989) direct method because it does not rest on fragile assumptions. Specifically, multifractal DFA can detrend series only with polynomial trends and is not as effective for sinusoidal trends at a variety of frequencies (Bashan et al., 2008; Ignaccolo et al., 2010). Furthermore, the validity of estimating both *α* and *f* (*α*) from the same *H* is untested in many biological examples (Zamir, 2003). We prefer Chhabra and Jensen’s (1989) method exactly because it estimates *α* and *f*(*α*) independently, providing an estimate of multifractal-spectrum width without the risk of amplifying any errors in multifractal DFA’s double use of the same *H*_fGn_ estimates.

This Chhabra and Jensen’s method (1989) samples non-negative series *u* (*t*) at progressively larger scales such that proportion of signal *P_i_* (*L*) falling within the *i^th^* bin of scale *L* is

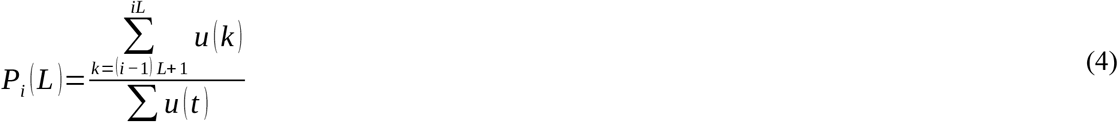

As *L* increases, *P_i_* (*L*) represents progressively larger proportion of *u* (*t*),

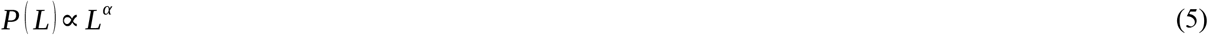

suggesting growth of proportion according to one ‘singularity’ strength *α*. *P* (*L*) exhibits multifractal dynamics when it grows heterogeneously across timescales *L* according to multiple potential fractional singularity strengths, such that

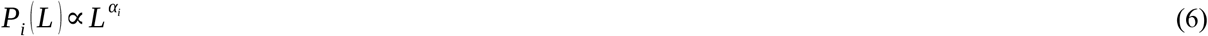

whereby each *i^th^* bin may show a distinct relationship of *P* (*L*) with *L*. The spectrum of singularities is itself the multifractal spectrum, and its width *Δ α*(*α_max_ – α_min_*) indicates the heterogeneity of these relationships between proportion and timescale (Halsey et al., 1986; Mandelbrot, 1999).

This method estimates *P* (*L*) for *N_L_* nonoverlapping bins of *L*-sizes and accentuates higher or lower *P* (*L*) by using parameter *q* >1 and *q* < 1, respectively, as follows

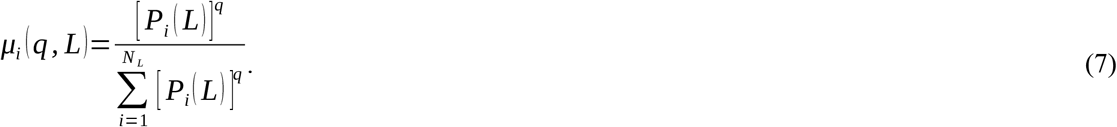

See *figure* 2.

*α* (*q*) is the singularity for *μ* (*q*)-weighted *P* (*L*) estimated by

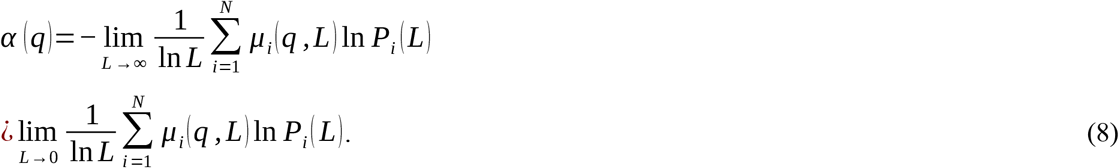

Estimates *α* (*q*) belong to the multifractal spectrum if Shannon entropy of *μ* (*q, l*) scales with *L* according to a dimension *f* (*q*), where

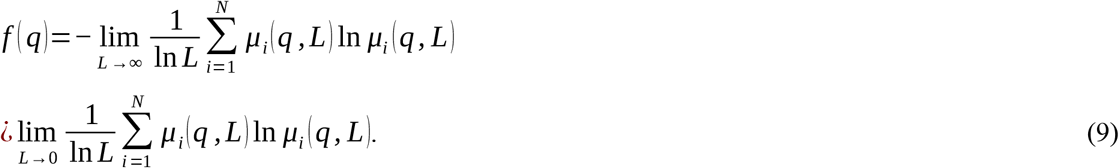

For *q* generating scaling relationships (Eqs. 8 and 9) with correlation coefficient, *r* > 0.9, the parametric curve (*α* (*q*), *f* (*q*)) or (*α, f* (*α*)) constitutes the multifractal spectrum with width *Δα* = *α_max_–α_min_. Δα* was estimated for each segment of CoM spatial-displacement series and CoP planar-displacement series (Figs. 2I and 2J).

To identify if nonzero *Δα* reflected nonlinear temporal correlations, *Δα* of each original series, was compared to *Δα* of 32 surrogate series obtained using Iterated Amplitude Adjusted Fourier Transformation (IAAFT) (Ihlen, 2012; Mandic et al., 2008). IAAFT generates surrogates with randomized phase ordering of the series’ spectral amplitudes but preserved linear temporal correlations. When *Δα* of the original series exceeds 1 95% confidence interval from *Δ α* of IAAFT series (i.e., *P* < 0.05), the original series is considered to have nonzero nonlinear correlations, easily quantifiable using the one-sample *t*-statistic (henceforth, CoP-*t*_MF_) comparing *Δ α* of the original series to that of the surrogates.

Finally, across all data (15 participants × 6 conditions/participant × 3 trials/condition × 12 segments/trial = 3240 segments), 3232/3240 or 99.75% CoM segments and 2935/3240 or 90.59% CoP segments exhibited multifractal spectrum that were significantly wider for the corresponding IAAFT surrogates (e.g., Fig. 3).

**Figure 3.**
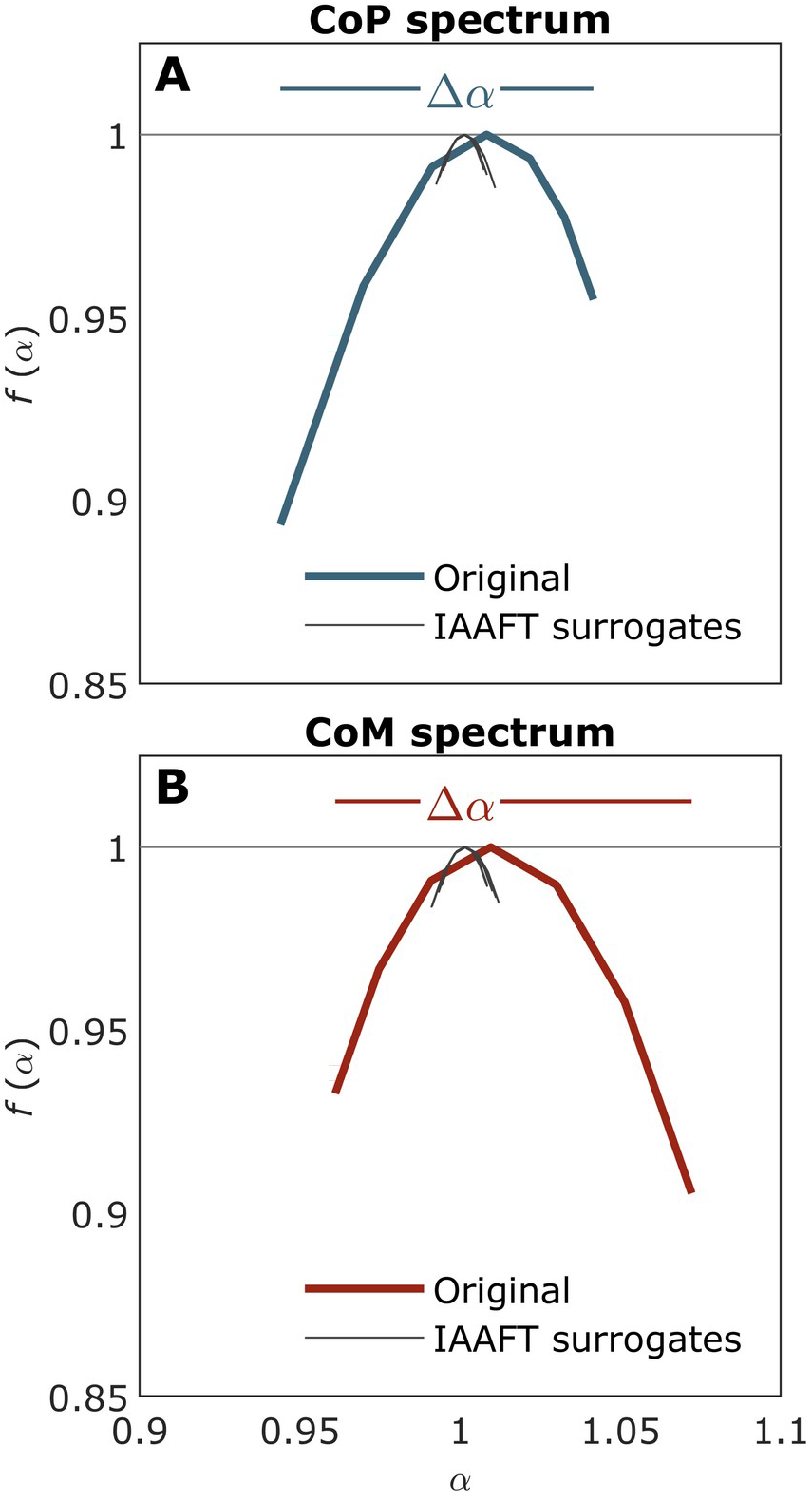
Multifractal spectra (*α, f*(*α*)) for the representative CoM spatial Euclidean series (**A**), and CoP planar Euclidean series (**B**), from Fig. 2.

### 2.6. Vector autoregression analysis

Vector autoregression captures linear interdependencies amongst concurrent series and here modeled intrapostural effects of CoP-*SD*, CoP-*H*_fGn_, and CoM-*H*_fGn_ in one segment on CoP-*SD*, CoP-*H*_fGn_, and CoM-*H*_gGn_ in subsequent segments (Hypotheses 2-4), and effects of CoP-*SD*, CoP-*H*_fGn_, and CoM-*t*_MF_ in one segment on CoP-*SD*, CoP-*H*_fGn_, and CoM-*t*_MF_ in subsequent segments (Hypothesis-5; Fig. 4). Vector autoregression describes each variable based on its own lagged value and that of each other variable. Lag is ideally increased until the residuals appear independently and are distributed identically (Sims, 1980).

**Figure 4.**
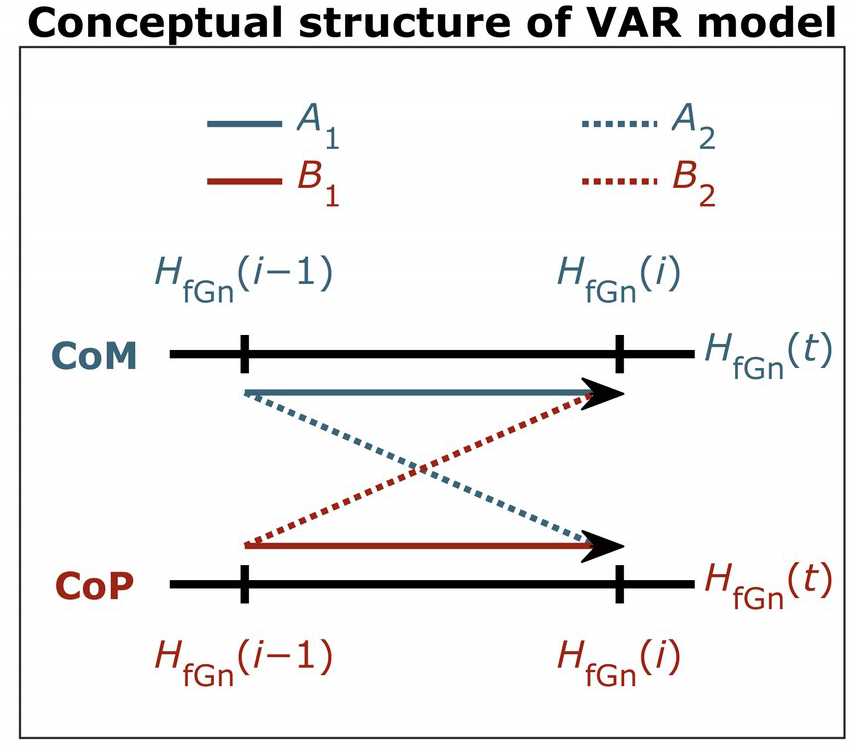
An overview of vector autoregressive (VAR) analysis. VAR analysis was used to model the diffusion of fractal fluctuations across the body, as a time series of segment-by-segment values of CoM-*H*_fGn_, CoP-*SD*, and CoP-*H*_fGn_. (Hypotheses-1–4), as well as CoP-*SD*, CoP-*H*_fGn_, and CoP-*t*_MF_ (Hypothesis-5). Black arrows indicate the effects of *H*_fGn_ in the previous segment on *H*_fGn_ in the current segment.

Vector autoregression allows forecasting unique effects of endogenous variables on later values of each other through impulse-response functions (IRFs). Impulse-response functions evaluate the relationship between *f* (*t*) and *g* (*t* + *τ*), or between *g* (*t*) and *f* (*t*+*τ*), where *τ* is a whole number corresponding to a segment within a trial. Provided vector-autoregression residuals are independent and identically distributed (i.i.d.), orthongonalizing these residuals allows simulating an ‘impulse’ to the system by adding 1*SEM* to any single variable, and using vector-autoregression coefficients to describe propagation of later ‘responses’ across all endogenous variables. The impulse-response function describes how an impulse in one series changes later predicted values in a different time series (Hatemi-J, 2004; Lutkepohl, 2007). All vector autoregression models converged with lag 1, with residuals passing all tests for i.i.d. status. We performed vector autoregression analysis using the function VAR() from the *vars* package (Pfaff et al., 2018) in *RStudio* (Team, 2013).

### 2.7. Statistical analysis of pairwise impulse-response functions

#### 2.7.1. Predictors

Mixed-effect regression models (Singer and Willett, 2003) treated impulse-response functions between each pair of postural descriptors as the dependent measure, testing the main effects of ‘Eyes Closed’ (EC; equaling 1 for the eyes-closed condition and 0 for all other conditions), and ‘Comfortable Viewing Distance’ (CVD; equaling 1 for 50-cm viewing condition and 0 for all other conditions), and ‘Distance From Comfortable Viewing Distance’ (DF_CVD: the absolute distance of viewing conditions from 50 for viewing conditions, i.e., 25, 0, 85, 170, and 255, as well as 0 for the eyes-closed condition), as well as each of the interaction of each of these main effects with the full-factorial family of interactions Trial × Segment × Impulse × Response. Impulse and Response served as class variables encoding the different descriptors (CoM-*H*_fGn_, CoP-*SD*, and CoP-*H*_fGn_ for Hypotheses-1–4; and including CoP-*t*_MF_ for Hypothesis 5; in both cases, CoM-*H*_fGn_ served as the baseline level for both Impulse and Response) serving as impulse and as response variables, respectively.

We thus used two mixed-effect regression models of the impulse-response functions: the first with the first set of impulse-response functions and the second with both sets of impulse-response functions (i.e., with and without multifractal contributions, respectively). In both cases, CoM-*H*_fGn_ served as the baseline level for both Impulse and Response. In the second regression model, we distinguished CoP-*H*_fGn_ as values of Impulse and CoP-*SD* as values of Response specifically to CoP-*t*_MF_ between the two vector autoregressions. (‘*’ after *H*_fGn_ and *SD* indicate the impulse-response functions from the second vector autoregression). The Impulse × Response terms highlighted significant differences of specific impulse-response functions from the global patterns. The description of results was made simpler by by the fact that the effects of multifractal nonlinearity outweighed all effects of monofractal evidence, as the second model showed no effects among the impulse-response functions from the first vector autoregression.

Each model was fit using the function lmer() from the *lme4* package (Bates et al., 2015) in *RStudio* (Team, 2013), with only a random-effect intercept for each participant. In the second model, a unique intercept was used for each participant in each vector autoregression to control for individual differences in impulse-response functions within each vector autoregression. Pearson’s correlation of residuals with model predictions was near zero for each of the two models (*r* = 0.0027, *p* = 0.679; and *r* = 0.0083, *p* = 0.067).

#### 2.7.2 Mixed-effect regression modeling for Hypotheses 1-4 and then Hypothesis 5 addressed impulse-response functions for only one vector-autoregressive model and both vector-autoregressive models, respectively

Hypotheses 1–4 involved modeling the impulse-response functions from a vector-autoregressive model, including only CoM-*H*_fGn_, CoP-*SD*, and CoP-*H*_fGn_ (Dataset S1). Including CoP-*t*_MF_ as a fourth variable led to a vector-autoregressive model that failed to converge with i.i.d. residuals. Therefore, because CoM-*H*_fGn_ ultimately exhibited little influence on the other variables, we omitted it from the second vector-autoregressive model, which included only CoP-*SD*, CoP-*H*_fGn_, and CoP-*t*_MF_. Hypothesis-5 involved modeling the cumulative set of impulse-response functions from both vector-autoregressive models (Dataset S2). This modeling choice reflects our interest in developing Hypothesis-5 as an elaboration of Hypothesis-1–4 without amplifying Type-I error. Hypothesis-5 requires additional impulse-response functions that Hypotheses-1–4 does not. Pooling all impulseresponse functions allowed testing Hypothesis-5 above and beyond the earlier-hypothesized structure instead of testing it separately in parallel.

#### 2.7.3. Orthogonal polynomials for segments within trial

Following a convention that we had developed in earlier work (Mangalam et al., 2020a, 2020b), orthogonal linear, quadratic, and cubic polynomials of Segment modeled how impulse-response functions changed over 999-sample segments within a trial. The cubic polynomial was used to capture the nonlinear decay of impulse-response functions across later segments. The interactions with Segment indicated changes in these effects with different third-order polynomial responses over subsequent trials. However, the cubic polynomial failed to improve model fit, so was omitted. The second regression model was much larger to model impulse-response functions from two separate vector autoregressions. The inclusion of multifractal nonlinearity within vector-autoregressive modeling had an enormous effect on reshaping the impulse-response functions. One of the consequences was the elaboration of the polynomial structure. The maximum-order polynomial for modeling across impulse-response values for each of 10 subsequent segments was 9, and the 9^th^-order polynomial significantly improved model fit (*χ*^2^(384) = 869.64, *p* < 0.0001). Because this 9^th^-order polynomial interacted only with *t*_MF_ as Impulse and as Response, we used 9^th^-order polynomials for the second model but not the first model. However, the second model omitted interactions of Trial with EC and DF_CVD because these interactions no longer improved model fit when testing Hypothesis-5—this failure of Trial interactions to improve model fit was equally true whether the second model included only 3^rd^- or all 9^th^-order polynomials, (*χ*^2^(256) = 164.04 and *χ*^2^(640) = 294.56, *P*s = 1.000).

#### 2.7.4 Addressing multiple comparisons by nesting hypotheses and testing new hypotheses while preserving data and model structure for earlier hypothesis

Despite the abundance of coefficients reported below, we stress that we report below the outcomes of only two regression models. Hypothesis-5 was nested both within the data and the model for Hypotheses-1–4— and not in a separate, new model. We inferred significance using *P*-values, with alpha set to the conventional 0.05 level in each model. Vector autoregression and impulse-response modeling appear here only to provide average measures, and while it is possible to estimate p-values for these VAR- and impulse-response-produced coefficients, they were not used here. Instead, differences among impulse-response model-generated coefficients were tested through a cumulative multiple-regression modeling procedure that retained, for Hypothesis-5, all previous structures for testing Hypotheses-1–4.

Such multiple regression allows simultaneously testing multiple predictors, precluding the need for addition correction (Williams, 1971). Furthermore, traditional corrections (e.g., Bonferroni) are apt when the dependent variable is correlated. However, because the test of Hypothesis-5 operates on both new impulse-response functions and old impulse-response functions appearing in the test of Hypotheses-1–4, the dependent variables for these distinct hypotheses cannot be correlated in any standard way. Retaining the impulse-response functions for Hypotheses-1–4 in the dataset while testing additional impulse-response functions ensured that the second model incorporated all of the structure of the first model.

## 3. Results

### 3.1. Regression modeling of impulse-response functions relating CoM-H_fGn_, CoP-SD, and CoP-H_fGn_

#### 3.1.1. Maintaining a quiet stance with eyes closed weakened intrapostural interactivity (Hypothesis-1)

The only significant effect in the eyes-closed condition on the intercept of the impulse-response functions was the one representing responses of CoP-*H*_fGn_ impulses (EC × Impulse(CoP-*H*_fGn_): *B* = 7.06×10^-3^, *P* = 2.88×10^-2^). This coefficient canceled out the negative responses of CoP-*H*_fGn_ impulses (Impulse(CoP-*H*_fGn_): *B* = −6.84×10^-3^, *P* = 1.73×10^-3^). All other significant effects in the regression refereed to polynomial changes in impulse-response functions (Tables 1 & 2), suggesting that the eyes-closed condition yielded impulse-response functions varying according to a polynomial centered on a zero mean.

**Table 1.** Significant baseline effects common to all conditions from mixed-effect regression modeling^a^ of impulse-response functions relating CoM-*H*_fGn_, CoP-*SD*, and CoP-*H*_fGn_ (see Section 3.1.1.).

| Effect | *B* | *SE* | *t* | *P*<sup>b</sup> |
| --- | --- | --- | --- | --- |
| poly(Segment,2)1 | $-5.25 \times 10^{-1}$ | $2.41 \times 10^{-1}$ | -2.18 | **$2.92 \times 10^{-2}$** |
| Impulse(CoP-$H_{fGn}$) | $-6.84 \times 10^{-3}$ | $2.18 \times 10^{-3}$ | -3.13 | **$1.73 \times 10^{-3}$** |
| Trial × poly(Segment,2)1 | $2.96 \times 10^{-1}$ | $1.11 \times 10^{-1}$ | 2.66 | **$7.84 \times 10^{-3}$** |
| poly(Segment,2)1 × Impulse(CoP-$H_{fGn}$) | $1.25 \times 10^0$ | $3.40 \times 10^{-1}$ | 3.67 | **$2.43 \times 10^{-4}$** |
| poly(Segment,2)1 × Response(CoP-$H_{fGn}$) | $8.67 \times 10^{-1}$ | $3.40 \times 10^{-1}$ | 2.55 | **$1.08 \times 10^{-2}$** |
| Impulse(CoP-$H_{fGn}$) × Response(CoP-$SD$) | $7.24 \times 10^{-3}$ | $3.10 \times 10^{-3}$ | 2.34 | **$1.91 \times 10^{-2}$** |
| Trial × poly(Segment,2)1 × Impulse(CoP-$H_{fGn}$) | $-5.47 \times 10^{-1}$ | $1.58 \times 10^{-1}$ | -3.47 | **$5.23 \times 10^{-4}$** |
| Trial × poly(Segment,2)1 × Response(CoP-$H_{fGn}$) | $-4.95 \times 10^{-1}$ | $1.58 \times 10^{-1}$ | -3.14 | **$1.68 \times 10^{-3}$** |
| poly(Segment,2)1 × Impulse(CoP-$H_{fGn}$) × Response(CoP-$H_{fGn}$) | $-1.15 \times 10^0$ | $4.81 \times 10^{-1}$ | -2.40 | **$1.65 \times 10^{-2}$** |
| poly(Segment,2)1 × Impulse(CoP-$H_{fGn}$) × Response(CoP-$SD$) | $-1.25 \times 10^0$ | $4.81 \times 10^{-1}$ | -2.60 | **$9.23 \times 10^{-3}$** |
| Trial × poly(Segment,2)1 × Impulse(CoP-$H_{fGn}$) × Response(CoP-$H_{fGn}$) | $5.73 \times 10^{-1}$ | $2.23 \times 10^{-1}$ | 2.57 | **$1.02 \times 10^{-2}$** |
| Trial × poly(Segment,2)1 × Impulse(CoP-$SD$) × Response(CoP-$H_{fGn}$) | $4.85 \times 10^{-1}$ | $2.23 \times 10^{-1}$ | 2.18 | **$2.94 \times 10^{-2}$** |
| Trial × poly(Segment,2)1 × Impulse(CoP-$H_{fGn}$) × Response(CoP-$SD$) | $5.43 \times 10^{-1}$ | $2.23 \times 10^{-1}$ | 2.44 | **$1.48 \times 10^{-2}$** |

<sup>a</sup>Fitted model: $IRF \sim (EC + CVD + DF\_CVD) * Trial * poly(segment,2) * Impulse * Response + (1 | Participant)$
<sup>b</sup>Boldfaced values indicate statistical significance at the the alpha level of 0.05.

**Table 2.** Significant effects of Eyes Closed (EC) from mixed-effect regression modeling^a^ of impulse-response functions relating CoM-*H*_fGn_, CoP-*SD*, and CoP-*H*_fGn_ (see Section 3.1.2.).

| Effect | *B* | *SE* | *t* | *p*<sup>b</sup> |
| --- | --- | --- | --- | --- |
| EC × poly(Segment,2)2 | $-9.57 \times 10^{-1}$ | $3.56 \times 10^{-1}$ | -2.69 | **$7.16 \times 10^{-3}$** |
| EC × Impulse(CoP-$H_{fGn}$) | $7.06 \times 10^{-3}$ | $3.23 \times 10^{-3}$ | 2.19 | **$2.88 \times 10^{-2}$** |
| EC × Trial × poly(Segment,2)2 | $5.46 \times 10^{-1}$ | $1.65 \times 10^{-1}$ | 3.31 | **$9.23 \times 10^{-4}$** |
| EC × poly(Segment,2)1 × Impulse(CoP-$H_{fGn}$) | $-1.95 \times 10^0$ | $5.03 \times 10^{-1}$ | -3.88 | **$1.04 \times 10^{-4}$** |
| EC × poly(Segment,2)2 × Impulse(CoP-$H_{fGn}$) | $1.63 \times 10^0$ | $5.03 \times 10^{-1}$ | 3.24 | **$1.21 \times 10^{-3}$** |
| EC × poly(Segment,2)2 × Impulse(CoP-$SD$) | $1.15 \times 10^0$ | $5.03 \times 10^{-1}$ | 2.28 | **$2.27 \times 10^{-2}$** |
| EC × Trial × poly(Segment,2)1 × Impulse(CoP-$H_{fGn}$) | $1.03 \times 10^0$ | $2.33 \times 10^{-1}$ | 4.43 | **$9.37 \times 10^{-6}$** |
| EC × Trial × poly(Segment,2)2 × Impulse(CoP-$H_{fGn}$) | $-1.05 \times 10^0$ | $2.33 \times 10^{-1}$ | -4.51 | **$6.51 \times 10^{-6}$** |
| EC × Trial × poly(Segment,2)2 × Impulse(CoP-$SD$) | $-6.69 \times 10^{-1}$ | $2.33 \times 10^{-1}$ | -2.87 | **$4.10 \times 10^{-3}$** |
| EC × Trial × poly(Segment,2)2 × Response(CoP-$H_{fGn}$) | $-5.25 \times 10^{-1}$ | $2.33 \times 10^{-1}$ | -2.25 | **$2.42 \times 10^{-2}$** |
| EC × Trial × poly(Segment,2)2 × Response(CoP-$SD$) | $-5.50 \times 10^{-1}$ | $2.33 \times 10^{-1}$ | -2.36 | **$1.83 \times 10^{-2}$** |
| EC × poly(Segment,2)1 × Impulse(CoP-$H_{fGn}$) × Response(CoP-$H_{fGn}$) | $1.72 \times 10^0$ | $7.12 \times 10^{-1}$ | 2.41 | **$1.58 \times 10^{-2}$** |
| EC × poly(Segment,2)2 × Impulse(CoP-$H_{fGn}$) × Response(CoP-$H_{fGn}$) | $-1.74 \times 10^0$ | $7.12 \times 10^{-1}$ | -2.45 | **$1.45 \times 10^{-2}$** |
| EC × poly(Segment,2)1 × Impulse(CoP-$H_{fGn}$) × Response(CoP-$SD$) | $1.97 \times 10^0$ | $7.12 \times 10^{-1}$ | 2.76 | **$5.72 \times 10^{-3}$** |
| EC × poly(Segment,2)2 × Impulse(CoP-$H_{fGn}$) × Response(CoP-$SD$) | $-1.66 \times 10^0$ | $7.12 \times 10^{-1}$ | -2.33 | **$2.00 \times 10^{-2}$** |
| EC × Trial × poly(Segment,2)1 × Impulse(CoP-$H_{fGn}$) × Response(CoP-$H_{fGn}$) | $-1.13 \times 10^0$ | $3.30 \times 10^{-1}$ | -3.42 | **$6.31 \times 10^{-4}$** |
| EC × Trial × poly(Segment,2)2 × Impulse(CoP-$H_{fGn}$) × Response(CoP-$H_{fGn}$) | $1.29 \times 10^0$ | $3.30 \times 10^{-1}$ | 3.90 | **$9.60 \times 10^{-5}$** |
| EC × Trial × poly(Segment,2)2 × Impulse(CoP-$SD$) × Response(CoP-$H_{fGn}$) | $7.26 \times 10^{-3}$ | $3.30 \times 10^{-1}$ | 2.20 | **$2.76 \times 10^{-2}$** |
| EC × Trial × poly(Segment,2)1 × Impulse(CoP-$H_{fGn}$) × Response(CoP-$SD$) | $-1.02 \times 10^0$ | $3.30 \times 10^{-1}$ | -3.11 | **$1.89 \times 10^{-3}$** |
| EC × Trial × poly(Segment,2)2 × Impulse(CoP-$H_{fGn}$) × Response(CoP-$SD$) | $1.06 \times 10^0$ | $3.30 \times 10^{-1}$ | 3.22 | **$1.30 \times 10^{-3}$** |
| EC × Trial × poly(Segment,2)2 × Impulse(CoP-$SD$) × Response(CoP-$SD$) | $6.77 \times 10^{-1}$ | $3.30 \times 10^{-1}$ | 2.05 | **$3.99 \times 10^{-2}$** |

<sup>a</sup>Fitted model: $IRF \sim (EC + CVD + DF\_CVD) * Trial * poly(segment,2) * Impulse * Response + (1|Participant)$
<sup>b</sup>Boldfaced values indicate statistical significance at the the alpha level of 0.05.

#### 3.1.2. CoM-H_fGn_ and CoP-H_fGn_ self-corrected from segment to segment within a trial but showed sparse effects on each other (Hypothesis-2)

In viewing conditions, increases in CoM-*H*_fGn_ and CoP-*H*_fGn_ predicted subsequent increases and reductions in alternation over segments (Figs. 5A and 5B). Visually fixating prompted monofractality to fall in and out of zero change or to cycle around zero change with negative and positive changes following each other, suggesting that viewing conditions prompted a sort of self-correcting maintenance of monofractality within CoM and CoP. The model did not yield significant impulse-response functions between CoM-*H*_fGn_ and CoP-*H*_fGn_ either (Figs. 5C and 5D): only the 220-cm viewing condition was accompanied by a CoM-*H*_fGn_ impulses that predicted CoP-*H*_fGn_ responses (Fig. 5C).

**Figure 5.**
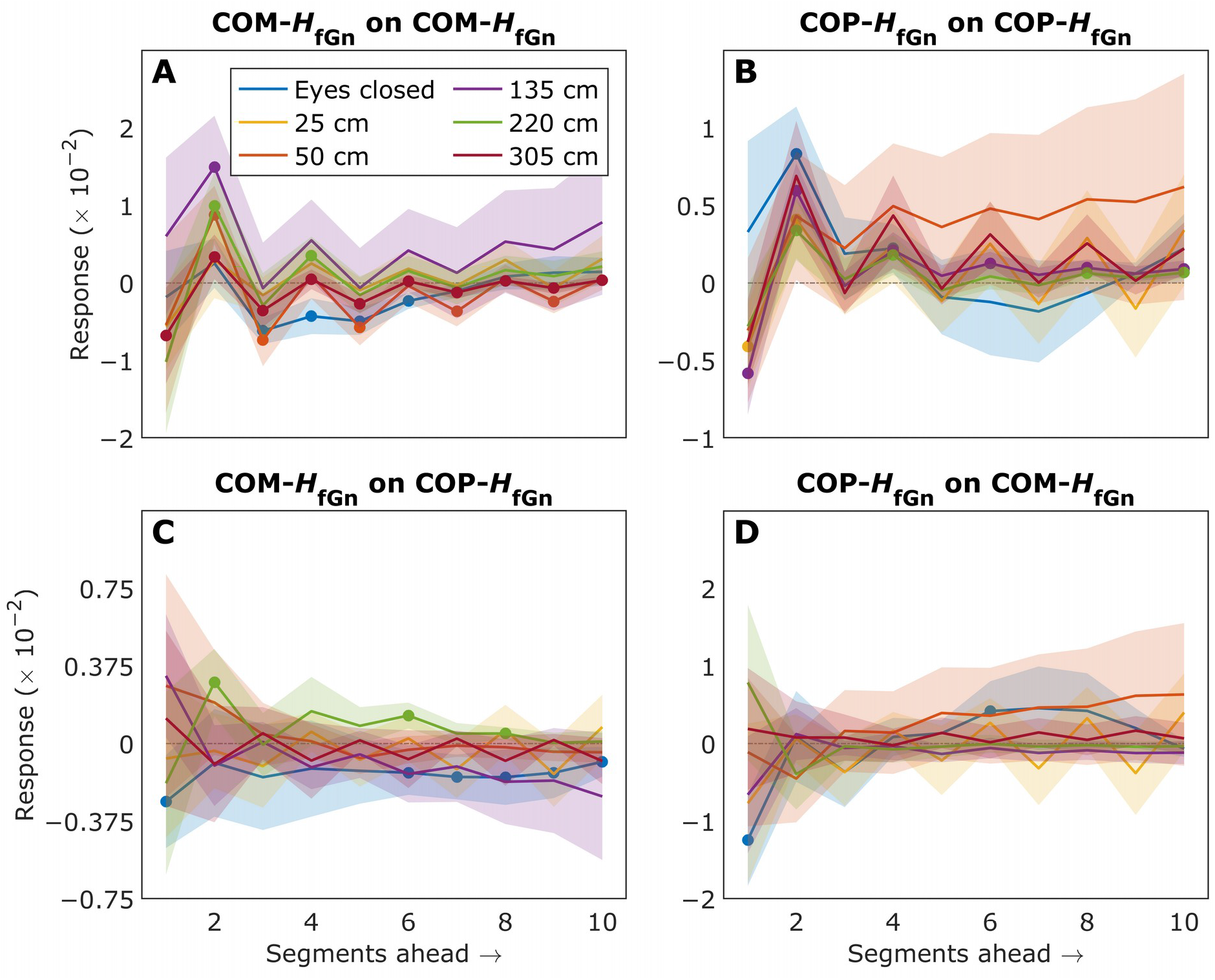
Impulse-response functions predicting the responses over ten segments ahead to an impulse in the current segment for each viewing condition. (**A**) CoM-*H*_fGn_ on CoM-*H*_fGn_. (**B**) CoP-*H*_fGn_ on CoP-*H*_fGn_. (**C**) CoM-*H*_fGn_ on CoP-*H*_fGn_.(**D**) CoP-*H*_fGn_ on CoM-*H*_fGn_. Shaded areas indicate *M*±1*SEM* of all trials across all participants (*N* = 15). Solid circles indicate statistically significant (*P* < 0.01) responses to an impulse in the *i*^th^ segment. The curves eventually approach zero, indicating that impulse-responses weakened over subsequent segments and eventually diminished completely.

#### 3.1.3. Increases in CoP-H_fGn_ and CoP-SD predicted subsequent reductions in each other across trials (Hypothesis-3)

In viewing conditions, CoP-*H*_fGn_ and CoP-*SD* impulses both predicted subsequent reductions in CoP-*SD* and CoP-*H*_fGn_, respectively (Fig. 6). These impulse-response functions remained robust with Trial, more so for the effects of CoP-*SD* impulses on CoP-*H*_fGn_ responses (Fig. 5B) than for the effects of CoP-*H*_fGn_ impulses on CoP-*SD* responses (Fig. 5A). 305-cm viewing condition failed to show a significant effect of CoP-*SD* impulses on CoP-*H*_fGn_ responses on only one trial. The 25-, 135-, and 220-cm viewing conditions each exhibited one, one, and two trials, respectively, that failed to show the effects of CoP-*H*_fGn_ impulses on CoP-*SD* responses.

**Figure 6.**
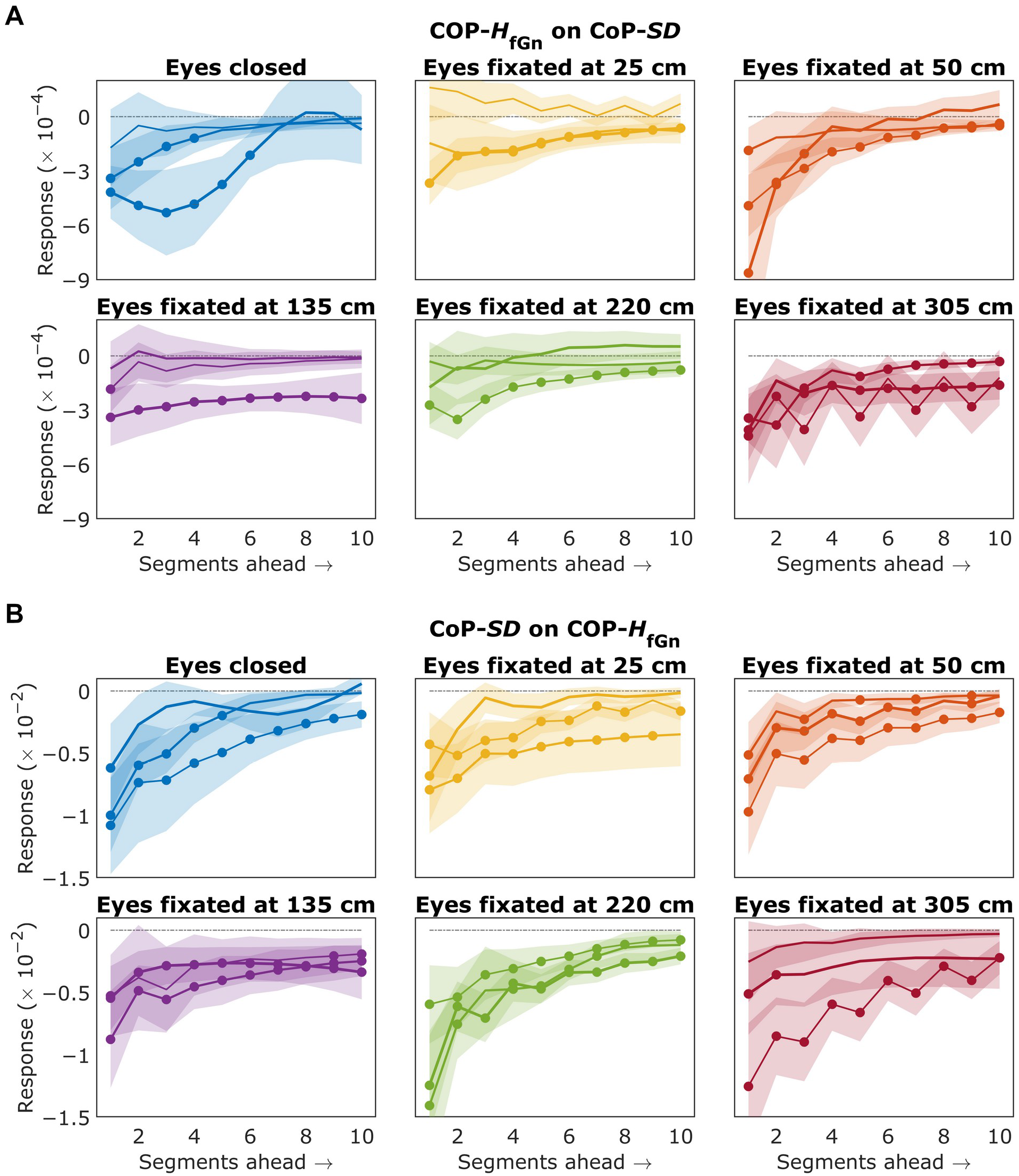
Impulse-response functions predicting the responses over ten segments ahead to an impulse in the current segment for each trial for each viewing condition. (**A**) CoP-*H*_fGn_ on CoP-*SD*. (**B**) CoP-*SD* on CoP-*H*_fGn_. Line widths encode trial order (thin: trial-1; medium: trial-2; thick: trial-3). Shaded areas indicate *M*±1*SEM* of trial averages across all participants (*N* = 15). Solid circles indicate statistically significant (*P* < 0.01) responses to an impulse in the *i*^th^ segment. The curves eventually approach zero, indicating that impulse-responses weakened over subsequent segments and eventually diminished completely.

#### 3.1.4. Impulse-response relationship between increases in CoP-H_fGn_ on subsequent reductions in CoP-SD was weakest for 50-cm viewing condition and became progressively stronger as viewing distance deviated from 50 cm (Hypothesis-4)

Of all viewing conditions, the 50-cm viewing condition prompted the least evidence that increases in CoP-*H*_fGn_ predicted subsequent reductions in CoP-*SD* (Table 3, Fig. 6A). For this comfortable viewing distance of 50 cm (i.e., Distance from Comfortable Viewing Distance, DF_CVD = 0), the intercept for impulse-response relating CoP-*H*_fGn_ impulses to CoP-*SD* responses was effectively zero. The effect of CoP-*H*_fGn_ impulses on CoP-*SD* responses (Impulse(CoP-*H*_fGn_) × Response(CoP-*SD*): *B* = 7.24×10^-3^, *P* = 1.91×10^-2^) canceled out the general effect of CoP-*H*_fGn_ impulses (Impulse(CoP-*H*_fGn_): *B* = −6.84×10^-3^, *P* = 1.73×10^-3^). In fact, the only intrapostural interactivity that appeared for the comfortable viewing distance was that an increase in CoP-*SD* predicted a subsequent increase in CoM-*H*_fGn_ (CVD *×* Impulse(CoP-*SD*): *B* = 9.30×10^-3^, *P* = 3.98×10^-3^). The effects of CoP-*SD* impulses on subsequent CoP-*H*_fGn_ and CoP-*SD* responses canceled out this positive effect (CVD *×* Impulse(CoP-*SD*) × Response(CoP-*H*_fGn_): *B* = −1.03×10^-2^, *P* = 2.36×10^-2^; CVD *×* Impulse(CoP-*SD*) *×* Response(CoP-*SD*): *B* = −1.00×10^-2^, *P* = 2.85×10^-2^, Fig. 6B). The 50-cm viewing distance resembled the eyes-closed case in that it carried many of the polynomial terms, suggesting impulse-response functions that varied nonlinearly around a zero mean.

**Table 3.** Significant effects of Comfortable Viewing Distance (CVD) from mixed-effect regression modeling^a^ of impulse-response functions relating CoM-*H*_fGn_, CoP-*SD*, and CoP-*H*_fGn_ (see Section 3.1.3.).

| Effect | *B* | *SE* | *t* | *P*<sup>b</sup> |
| --- | --- | --- | --- | --- |
| CVD × poly(Segment,2)1 | 1.38×10<sup>0</sup> | 3.56×10<sup>-1</sup> | 3.88 | **1.05×10<sup>-4</sup>** |
| CVD × poly(Segment,2)2 | -9.42×10<sup>-1</sup> | 3.56×10<sup>-1</sup> | -2.65 | **8.14×10<sup>-3</sup>** |
| CVD × Impulse(CoP-$SD$) | 9.30×10<sup>-3</sup> | 3.23×10<sup>-3</sup> | 2.88 | **3.98×10<sup>-3</sup>** |
| CVD × Trial × poly(Segment,2)1 | -7.14×10<sup>-1</sup> | 1.65×10<sup>-1</sup> | -4.33 | **1.47×10<sup>-5</sup>** |
| CVD × Trial × poly(Segment,2)2 | 4.68×10<sup>-1</sup> | 1.65×10<sup>-1</sup> | 2.84 | **4.51×10<sup>-3</sup>** |
| CVD × Trial × Impulse(CoP-$H_{fGn}$) | 4.45×10<sup>-3</sup> | 1.49×10<sup>-3</sup> | 2.98 | **2.91×10<sup>-3</sup>** |
| CVD × poly(Segment,2)1 × Impulse(CoP-$H_{fGn}$) | -2.76×10<sup>0</sup> | 5.03×10<sup>-1</sup> | -5.49 | **4.03×10<sup>-8</sup>** |
| CVD × poly(Segment,2)2 × Impulse(CoP-$H_{fGn}$) | 1.02×10<sup>0</sup> | 5.03×10<sup>-1</sup> | 2.02 | **4.33×10<sup>-2</sup>** |
| CVD × poly(Segment,2)1 × Impulse(CoP-$SD$) | -1.92×10<sup>0</sup> | 5.03×10<sup>-1</sup> | -3.81 | **1.38×10<sup>-4</sup>** |
| CVD × poly(Segment,2)2 × Impulse(CoP-$SD$) | 1.13×10<sup>0</sup> | 5.03×10<sup>-1</sup> | 2.24 | **2.49×10<sup>-2</sup>** |
| CVD × poly(Segment,2)1 × Response(CoP-$H_{fGn}$) | -1.37×10<sup>0</sup> | 5.03×10<sup>-1</sup> | -2.71 | **6.69×10<sup>-3</sup>** |
| CVD × poly(Segment,2)1 × Response(CoP-$SD$) | -1.36×10<sup>0</sup> | 5.03×10<sup>-1</sup> | -2.70 | **6.98×10<sup>-3</sup>** |
| CVD × Impulse(CoP-$SD$) × Response(CoP-$H_{fGn}$) | -1.03×10<sup>-2</sup> | 4.57×10<sup>-3</sup> | -2.26 | **2.36×10<sup>-2</sup>** |
| CVD × Impulse(CoP-$SD$) × Response(CoP-$SD$) | -1.00×10<sup>-2</sup> | 4.57×10<sup>-3</sup> | -2.19 | **2.85×10<sup>-2</sup>** |
| CVD × Trial × poly(Segment,2)1 × Impulse(CoP-$H_{fGn}$) | 1.52×10<sup>0</sup> | 2.33×10<sup>-1</sup> | 6.52 | **7.05×10<sup>-11</sup>** |
| CVD × Trial × poly(Segment,2)2 × Impulse(CoP-$H_{fGn}$) | -4.78×10<sup>-1</sup> | 2.33×10<sup>-1</sup> | -2.05 | **4.01×10<sup>-2</sup>** |
| CVD × Trial × poly(Segment,2)1 × Impulse(CoP-$SD$) | 8.64×10<sup>-1</sup> | 2.33×10<sup>-1</sup> | 3.71 | **2.09×10<sup>-4</sup>** |
| CVD × Trial × poly(Segment,2)2 × Impulse(CoP-$SD$) | -4.81×10<sup>-1</sup> | 2.33×10<sup>-1</sup> | -2.07 | **3.89×10<sup>-2</sup>** |
| CVD × Trial × poly(Segment,2)1 × Response(CoP-$H_{fGn}$) | 6.67×10<sup>-1</sup> | 2.33×10<sup>-1</sup> | 2.86 | **4.20×10<sup>-3</sup>** |
| CVD × Trial × poly(Segment,2)1 × Response(CoP-$SD$) | 6.93×10<sup>-1</sup> | 2.33×10<sup>-1</sup> | 2.97 | **2.95×10<sup>-3</sup>** |
| CVD × Trial × poly(Segment,2)2 × Response(CoP-$SD$) | -4.61×10<sup>-1</sup> | 2.33×10<sup>-1</sup> | -1.98 | **4.81×10<sup>-2</sup>** |
| CVD × Trial × Impulse(CoP-$H_{fGn}$) × Response(CoP-$SD$) | -4.22×10<sup>-3</sup> | 2.11×10<sup>-3</sup> | -2.00 | **4.58×10<sup>-2</sup>** |
| CVD × poly(Segment,2)1 × Impulse(CoP-$H_{fGn}$) × Response(CoP-$H_{fGn}$) | 1.69×10<sup>0</sup> | 7.12×10<sup>-1</sup> | 2.38 | **1.75×10<sup>-2</sup>** |
| CVD × poly(Segment,2)1 × Impulse(CoP-$SD$) × Response(CoP-$H_{fGn}$) | 2.08×10<sup>0</sup> | 7.12×10<sup>-1</sup> | 2.93 | **3.44×10<sup>-3</sup>** |
| CVD × poly(Segment,2)1 × Impulse(CoP-$H_{fGn}$) × Response(CoP-$SD$) | 2.76×10<sup>0</sup> | 7.12×10<sup>-1</sup> | 3.88 | **1.06×10<sup>-4</sup>** |
| CVD × poly(Segment,2)1 × Impulse(CoP-$SD$) × Response(CoP-$SD$) | 1.92×10<sup>0</sup> | 7.12×10<sup>-1</sup> | 2.69 | **7.14×10<sup>-3</sup>** |
| CVD × Trial × poly(Segment,2)1 × Impulse(CoP-$H_{fGn}$) × Response(CoP-$H_{fGn}$) | -8.49×10<sup>-1</sup> | 3.29×10<sup>-1</sup> | -2.58 | **9.97×10<sup>-3</sup>** |
| CVD × Trial × poly(Segment,2)1 × Impulse(CoP-$SD$) × Response(CoP-$H_{fGn}$) | -8.88×10<sup>-1</sup> | 3.29×10<sup>-1</sup> | -2.69 | **7.06×10<sup>-3</sup>** |
| CVD × Trial × poly(Segment,2)1 × Impulse(CoP-$H_{fGn}$) × Response(CoP-$SD$) | -1.50×10<sup>0</sup> | 3.29×10<sup>-1</sup> | -4.55 | **5.43×10<sup>-3</sup>** |
| CVD × Trial × poly(Segment,2)1 × Impulse(CoP-$SD$) × Response(CoP-$H_{fGn}$) | -8.56×10<sup>-1</sup> | 3.29×10<sup>-1</sup> | -2.60 | **9.41×10<sup>-3</sup>** |

<sup>a</sup>Fitted model: $IRF \sim (EC + CVD + DF\_CVD) * Trial * poly(segment,2) * Impulse * Response + (1|Participant)$
<sup>b</sup>Boldfaced values indicate statistical significance at the the alpha level of 0.05.

For all other viewing distances, an increase in CoP-*H*_fGn_ predicted a subsequent reduction in CoP-*SD*, and it was a larger reduction as viewing distance deviated in either direction (DF_CVD × Impulse(CoP-*H*_fGn_): *B* = 5.03×10^-5^, *P* = 2.36×10^-4^; DF_CVD × Impulse(CoP-*H*_fGn_) × Response(CoP-*SD*): *B* = −5.33×10^-5^, *P* = 5.88×10^-3^, Table 4, Fig. 6A). This reduction in CoP-*SD* following an increase in CoP-*H*_fGn_ became progressively larger across trials (DF_CVD × Trial × Impulse(CoP-*H*_fGn_): *B* = −1.43×10^-5^, *P* = 2.36×10^-2^). For instance, the reduction in CoP-*SD* following an increase in CoP-*H*_fGn_ ranged from −3.83×10^-5^ to −1.13×10^-2^, for 25-cm viewing condition on Trial-1 to 255 cm on Trial-3, respectively (Table 1).

**Table 4.** Significant effects of Distance from Comfortable Viewing Distance (DF_CVD) from mixed-effect regression modeling^a^ of impulse-response functions relating CoM-*H*_fGn_, CoP-*SD*, and CoP-*H*_fGn_ (see Section 3.1.4.).

| Effect | *B* | *SE* | *t* | *P*<sup>b</sup> |
| --- | --- | --- | --- | --- |
| *Effects of Distance from Comfortable Viewing Distance (DF_CVD)* |  |  |  |  |
| DF_CVD × Impulse(CoP-$H_{fGn}$) | $5.03 \times 10^{-5}$ | $1.37 \times 10^{-5}$ | 3.68 | **$2.36 \times 10^{-4}$** |
| DF_CVD × Trial × Impulse(CoP-$H_{fGn}$) | $-1.43 \times 10^{-5}$ | $6.34 \times 10^{-6}$ | -2.26 | **$2.36 \times 10^{-2}$** |
| DF_CVD × poly(Segment,2)1 × Impulse(CoP-$H_{fGn}$) | $-4.91 \times 10^{-3}$ | $2.13 \times 10^{-3}$ | -2.30 | **$2.15 \times 10^{-2}$** |
| DF_CVD × Impulse(CoP-$H_{fGn}$) × Response(CoP-$H_{fGn}$) | $-5.21 \times 10^{-5}$ | $1.94 \times 10^{-5}$ | -2.69 | **$7.14 \times 10^{-3}$** |
| DF_CVD × Impulse(CoP-$H_{fGn}$) × Response(CoP-$SD$) | $-5.33 \times 10^{-5}$ | $1.94 \times 10^{-5}$ | -2.75 | **$5.88 \times 10^{-3}$** |
| DF_CVD × Trial × poly(Segment,2)1 × Response(CoP-$H_{fGn}$) | $1.97 \times 10^{-3}$ | $9.88 \times 10^{-4}$ | 2.00 | **$4.57 \times 10^{-2}$** |
| DF_CVD × poly(Segment,2)1 × Impulse(CoP-$SD$) × Response(CoP-$H_{fGn}$) | $6.09 \times 10^{-3}$ | $3.02 \times 10^{-3}$ | 2.02 | **$4.35 \times 10^{-2}$** |
| DF_CVD × Trial × poly(Segment,2)1 × Impulse(CoP-$SD$) × Response(CoP-$H_{fGn}$) | $-2.83 \times 10^{-3}$ | $1.40 \times 10^{-3}$ | -2.03 | **$4.28 \times 10^{-2}$** |

<sup>a</sup>Fitted model: $IRF \sim (EC + CVD + DF\_CVD) * Trial * poly(segment,2) * Impulse * Response + (1|Participant)$
<sup>b</sup>Boldfaced values indicate statistical significance at the the alpha level of 0.05.

### 3.2. Regression modeling of impulse-response functions incorporating CoP-t_MF_ along with the prior set of CoM-H_fGn_, CoP-H_fGn_, and CoP-SD

This second regression model pooled impulse-response functions from both vector autoregressions. Hence, before we detail specific effects in this model, it is noteworthy that this regression model found no effects within the impulse-response functions from the first model that excluded multifractal nonlinearity but only effects among the impulse-response functions from the vector autoregression that included multifractal nonlinearity. Hence, this initial summary point reflects that multifractal nonlinearity exerted much stronger effects on monofractal evidence or *SD* than either had exerted on the other.

We also note that the use of random-effect intercept for participant-by-participant vector autoregression will necessarily make the regression model structure diverge from the simpler plot of Segment-by-Segment means of the impulse-response functions. We regret this divergence for how it can make the results more challenging to digest. However, this random-effect intercept term’s usefulness addressed the fact that impulseresponse functions varied substantially across participants. We have elsewhere documented that the early values of impulse-response functions predict individual differences in perceptual responses and, consequently, in the use of available information (Mangalam et al., 2020a, 2020b). The regression model takes a deeper grasp of the impulse-response functions below the more idiosyncratic variation in raw magnitude with its 9^th^-order polynomial structure.

With the preceding caveat, we share Segment-by-Segment participant-wise averages (*N* = 15) of impulse-response functions in figures with individual panels for each condition (Figs. 7 and 8). The following results will speak to the model coefficients and illustrated through the model-predicted form of the impulseresponse functions in each case (Figs. 9–11).

**Figure 7.**
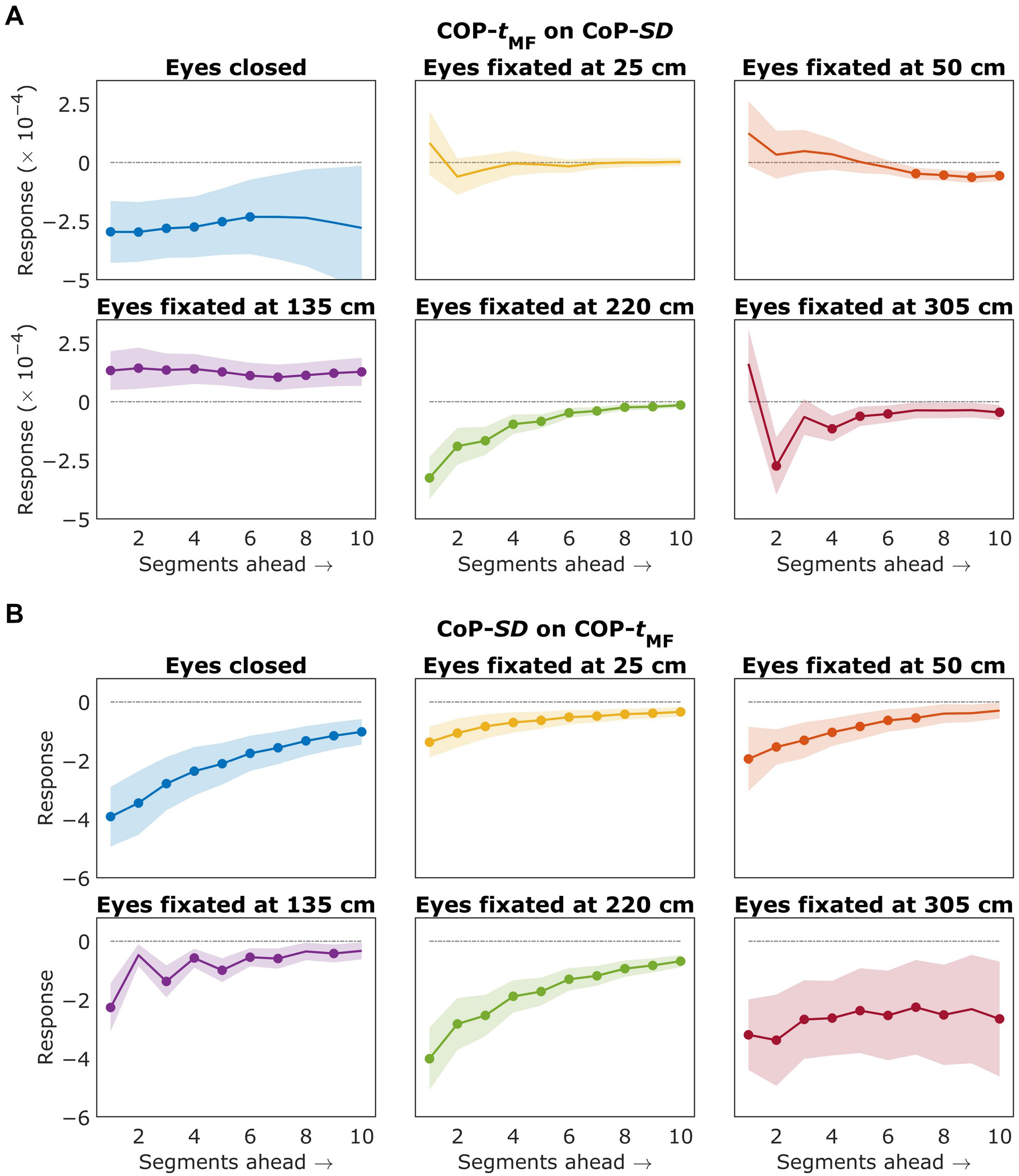
Impulse-response functions predicting the responses over ten segments ahead to an impulse in the current segment averaged across all trials for each condition. (**A**) CoP-*t*_MF_ on CoP-*SD*. In 50-cm viewing condition, a trial with outlier values was substituted by another trial for the same condition, for one participant. (**B**) CoP-*SD* on CoP-*t*_MF_. Shaded areas indicate *M*±1*SEM* of all trial across all participants (*N* = 45). Solid circles indicate statistically significant (*P* < 0.01) responses to an impulse in the *i*^th^ segment. The curves eventually approach zero, indicating that impulse-responses weakened over subsequent segments and eventually diminished completely.

**Figure 8.**
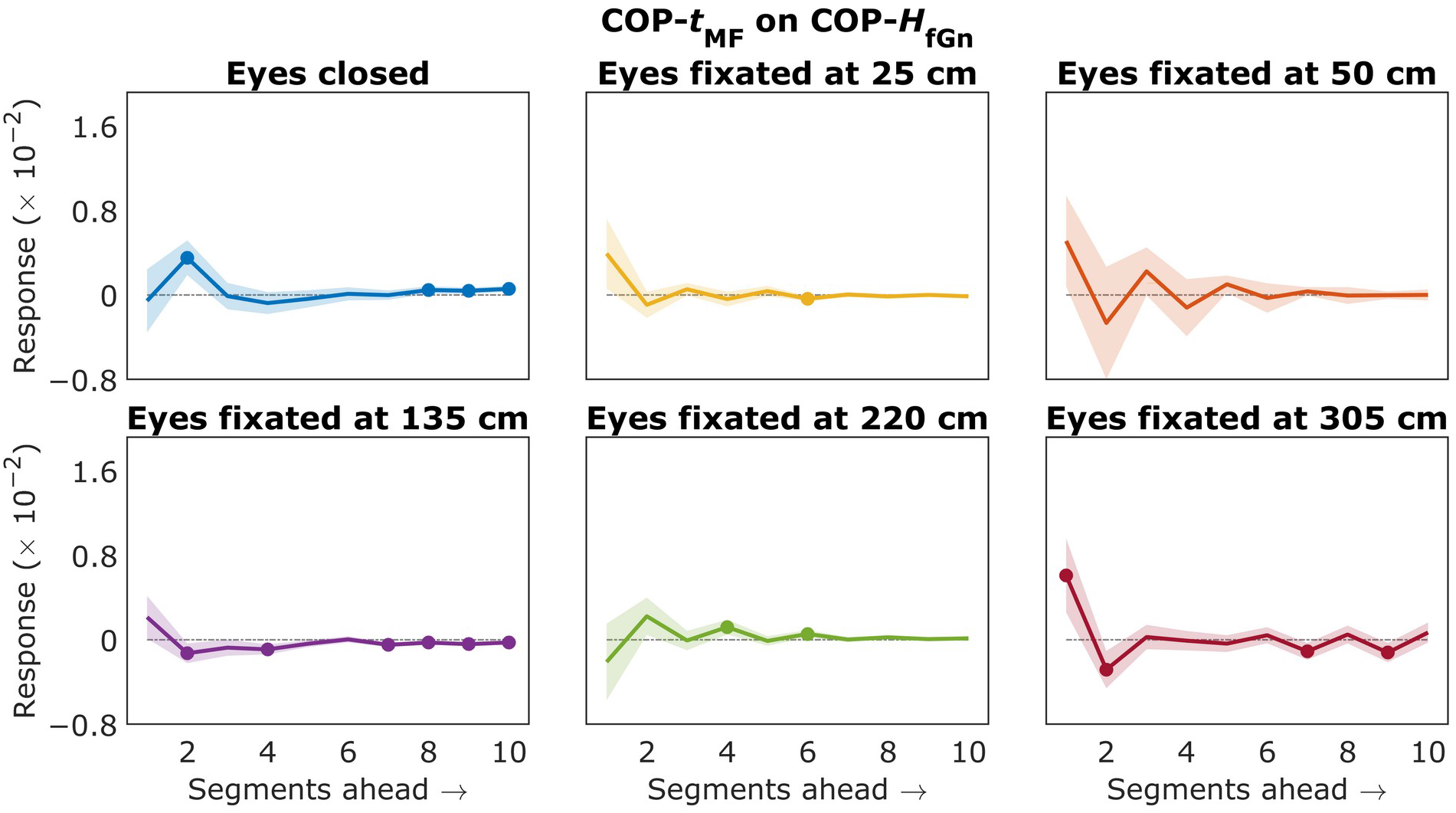
Impulse-response functions predicting the responses in CoP-*H*_fGn_ over ten segments ahead to an impulse in CoP-*t*_MF_ the current segment averaged across all trials for each condition. (**A**) CoP-*t*_MF_ on CoP-*SD*. (**B**) CoP-*SD* on CoP-*t*_MF_. In 305-cm viewing condition, a trial with outlier values was substituted by another trial for the same condition, for two participants. Shaded areas indicate *M*±1*SEM* of all trial across all participants (*N* = 45). Solid circles indicate statistically significant (*P* < 0.01) responses to an impulse in the *i*^th^ segment. The curves eventually approach zero, indicating that impulse-responses weakened over subsequent segments and eventually diminished completely.

**Figure 9.**
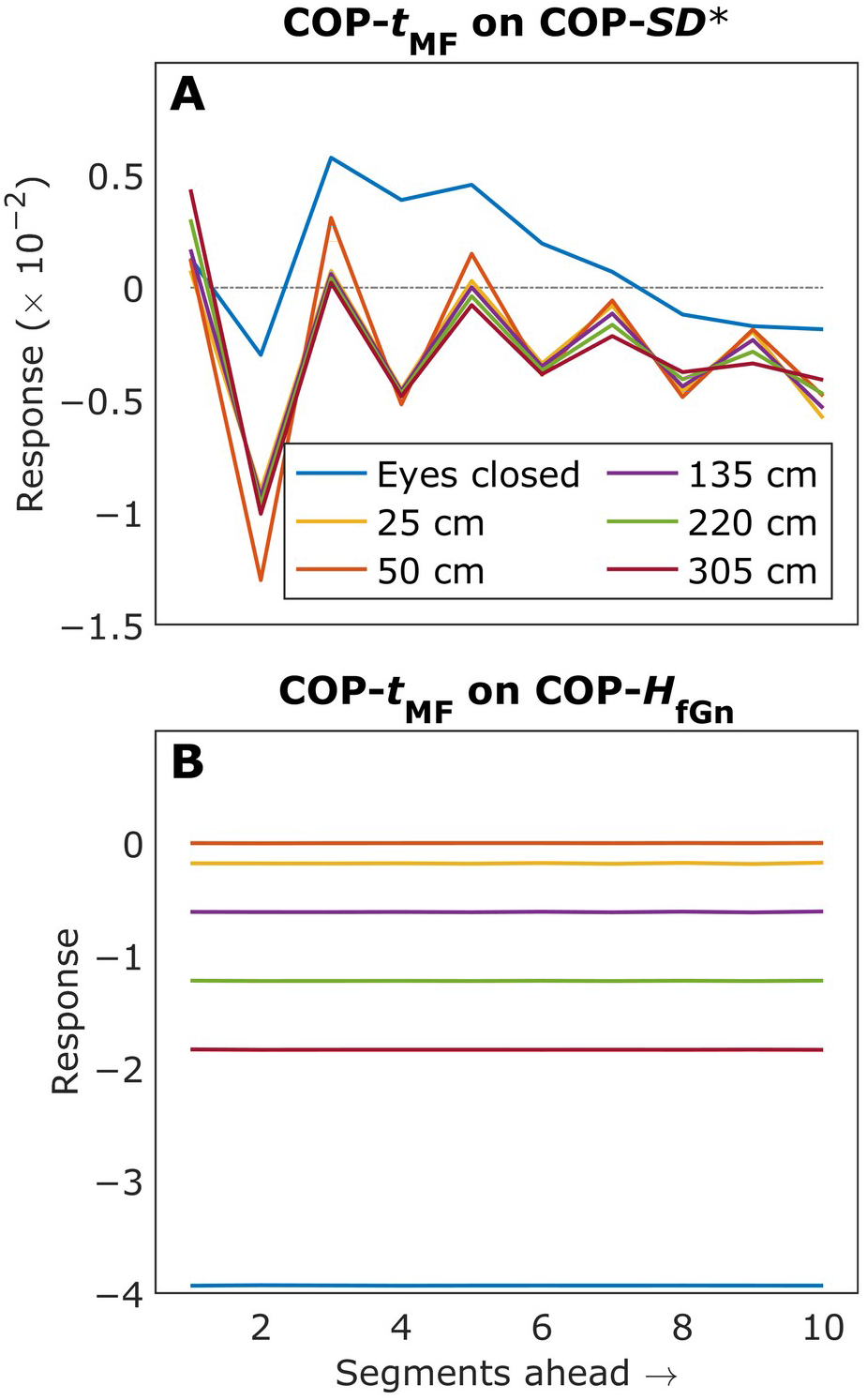
Impulse-response functions predicting the responses over ten segments ahead to an impulse in the current segment for each viewing condition. (**A**) CoP-*t*_MF_ on CoP-*SD*\*. (**B**) CoP-*t*_MF_ on on CoP-*H*_fGn_. ‘*’ after CoP-*SD* indicates the impulse-response functions from the second vector autoregression (see Section 2.7.1.).

**Figure 10.**
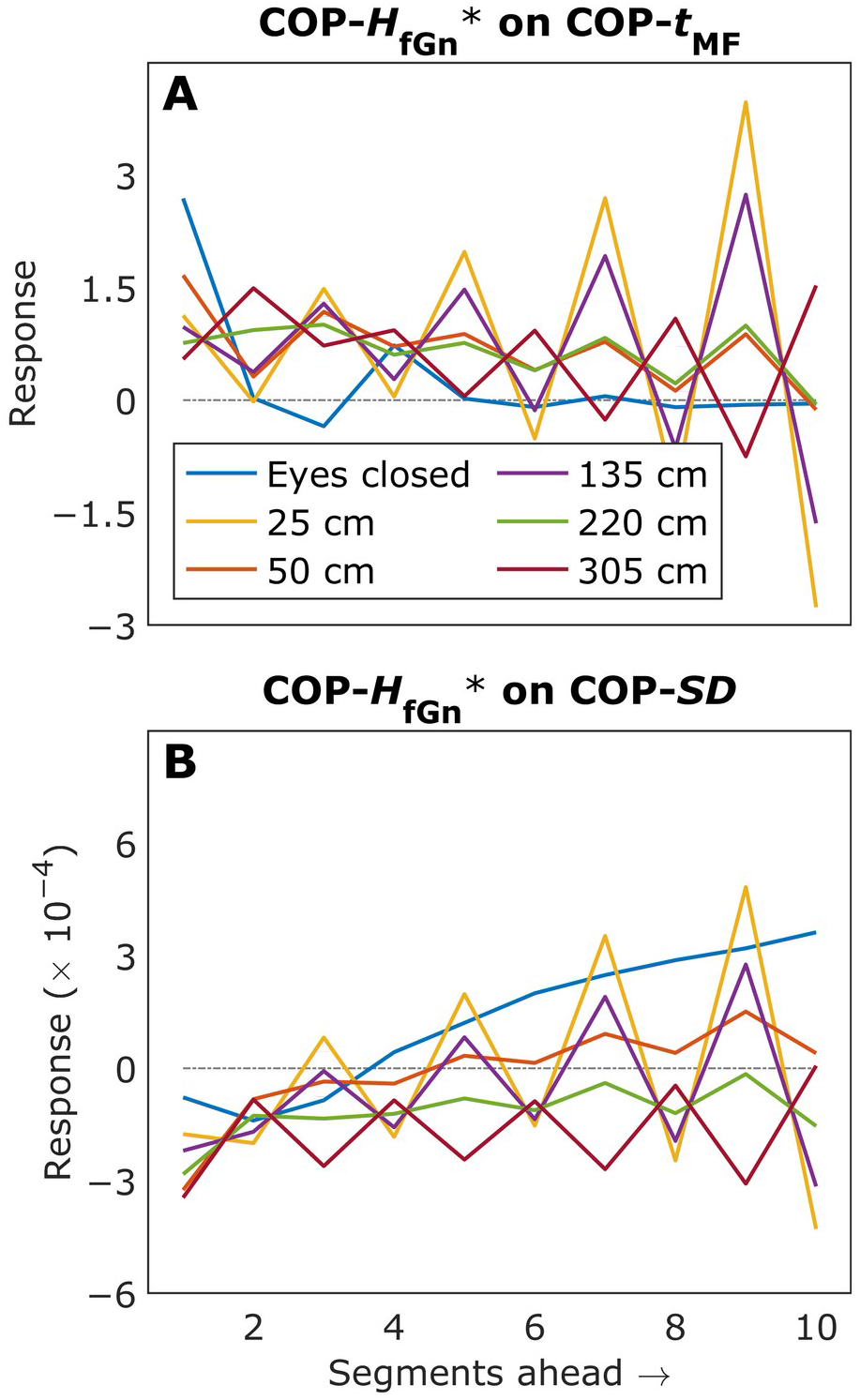
Impulse-response functions predicting the responses over ten segments ahead to an impulse in the current segment for each viewing condition. (**A**) CoP-*H*_fGn_* on CoP-*t*_MF_. (**B**) CoP-*H*_fGn_* on CoP-*SD*. ‘*’ after CoP-*H*_fGn_ indicates the impulse-response functions from the second vector autoregression (see Section 2.7.1.).

**Figure 11.**
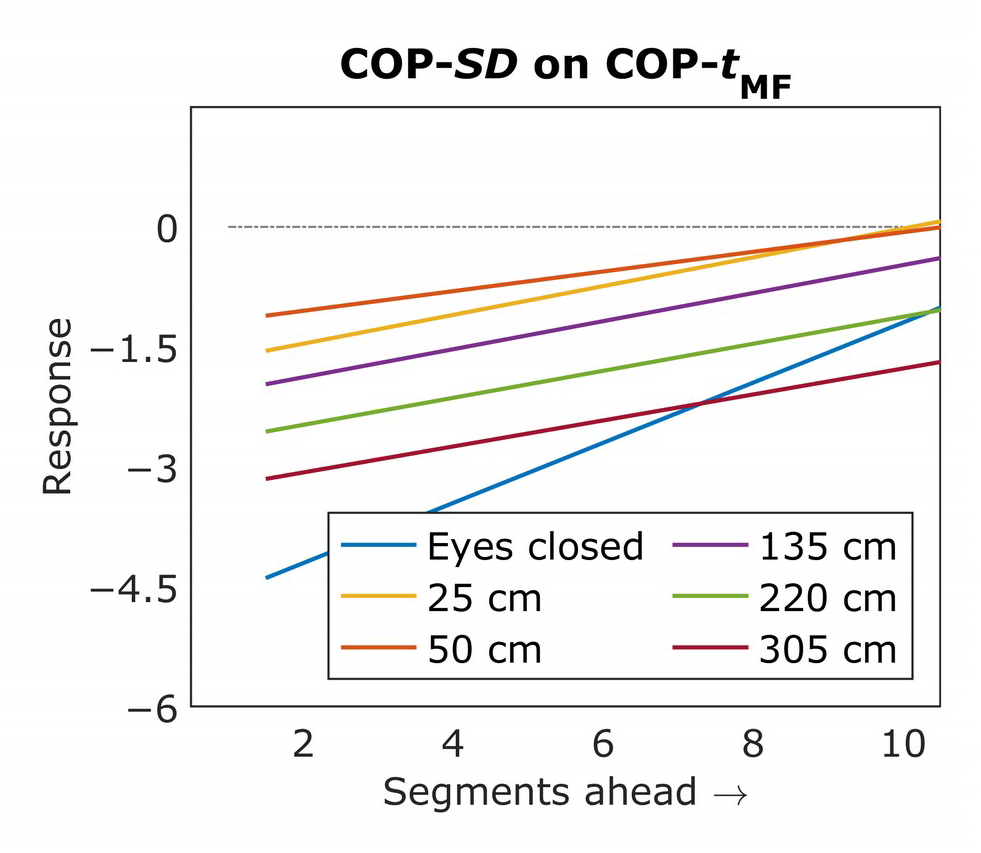
Impulse-response functions predicting the responses in CoP-*t*_MF_ over ten segments ahead to an impulse in CoP-*SD*.

#### 3.2.1. Increasing visual-precision demand led multifractal nonlinearity to predict larger subsequent decreases in SD of postural fluctuations (Hypothesis-5a)

In 50-cm viewing condition, increases in multifractal nonlinearity predicted oscillations in subsequent *SD* (Fig. 9A). After predicting an initial reduction in subsequent *SD* (2 segments ahead), increases in multifractal nonlinearity predicted a zig-zag pattern of subsequent changes reaching briefly back into positive values. That is, increasing multifractal nonlinearity predicted subsequent increases in *SD* (only 3 and 5 segments ahead). But this bobbling of subsequent changes in *SD* became progressively more negative, consistently predicting reductions in subsequent *SD*. This negative-then-briefly-positive-and-negative-again structure was specified by the 9^th^-order polynomials reflecting the impulse from CoP-*t*_MF_ (Impulse(CoP-*t*_MF_), *B* = 9.75×10^-1^, *P* = 1.12×10^-2^; 9^th^-order Segment × Impulse(CoP-*t*_MF_), polynomial coefficients from linear to nonic, *P*s < 0.05, Table 5; CVD × 9^th^-order Segment × Impulse(CoP-*t*_MF_), polynomial coefficients from cubic to nonic, *P*s < 0.05, Table 6) and the response from CoP-*SD* in 50-cm viewing condition (Response(CoP-*SD*\*), *B* = −9.78×10^-1^, *P* = 7.0010^-3^; 9^th^-order Segment all *P*s < 0.05, Table 5; Response(CoP-*SD*\*), polynomial coefficients from linear to nonic, *P*s < 0.05, Table 5; and CVD × 9^th^-order Segment × Response(CoP-*SD*) polynomial coefficients from linear to nonic, *P*s < 0.05, Table 6).

**Table 5.** Significant baseline effects common to all conditions from mixed-effect regression modeling^a^ of impulse-response functions incorporating CoP-*t*_MF_ along with the prior set of CoM-*H*_fGn_, CoP-*SD*, and CoP-*H*_fGn_ (see Section 3.2.1.). ‘*’ after *H*_fGn_ and *SD* indicate the impulse-response functions from the second vector autoregression.

| Effect | $B$ | $SE$ | $t$ | $P^b$ |
| --- | --- | --- | --- | --- |
| Impulse(CoP-$H_{fGn}$*) | $1.24 \times 10^0$ | $3.84 \times 10^{-1}$ | 3.22 | $1.28 \times 10^{-3}$ |
| poly(Segment, 9)1 × Impulse(CoP-$H_{fGn}$*) | $-1.77 \times 10^2$ | $8.05 \times 10^1$ | -2.20 | $2.80 \times 10^{-2}$ |
| poly(Segment, 9)3 × Impulse(CoP-$H_{fGn}$*) | $-1.68 \times 10^2$ | $8.05 \times 10^1$ | -2.09 | $3.70 \times 10^{-2}$ |
| poly(Segment, 9)5 × Impulse(CoP-$H_{fGn}$*) | $-2.17 \times 10^2$ | $8.05 \times 10^1$ | -2.70 | $7.03 \times 10^{-3}$ |
| poly(Segment, 9)7 × Impulse(CoP-$H_{fGn}$*) | $-2.22 \times 10^2$ | $8.05 \times 10^1$ | -2.75 | $5.92 \times 10^{-3}$ |
| poly(Segment, 9)9 × Impulse(CoP-$H_{fGn}$*) | $-2.72 \times 10^2$ | $8.05 \times 10^1$ | -3.38 | $7.33 \times 10^{-4}$ |
| Impulse(CoP-$H_{fGn}$*) × Response(CoP-$H_{fGn}$) | $-1.25 \times 10^0$ | $4.52 \times 10^{-1}$ | -2.77 | $5.58 \times 10^{-3}$ |
| poly(Segment, 9)5 × Impulse(CoP-$H_{fGn}$*) × Response(CoP-$H_{fGn}$) | $2.30 \times 10^2$ | $9.86 \times 10^1$ | 2.33 | $1.99 \times 10^{-2}$ |
| poly(Segment, 9)7 × Impulse(CoP-$H_{fGn}$*) × Response(CoP-$H_{fGn}$) | $2.29 \times 10^2$ | $9.86 \times 10^1$ | 2.33 | $2.01 \times 10^{-2}$ |
| poly(Segment, 9)9 × Impulse(CoP-$H_{fGn}$*) × Response(CoP-$H_{fGn}$) | $2.76 \times 10^2$ | $9.86 \times 10^1$ | 2.80 | $5.14 \times 10^{-3}$ |
| Impulse(CoP-$H_{fGn}$*) × Response(CoP-$SD$) | $-1.24 \times 10^0$ | $4.52 \times 10^{-1}$ | -2.74 | $6.16 \times 10^{-3}$ |
| poly(Segment, 9)5 × Impulse(CoP-$H_{fGn}$*) × Response(CoP-$SD$) | $2.17 \times 10^2$ | $9.86 \times 10^1$ | 2.20 | $2.78 \times 10^{-2}$ |
| poly(Segment, 9)7 × Impulse(CoP-$H_{fGn}$*) × Response(CoP-$SD$) | $2.22 \times 10^2$ | $9.86 \times 10^1$ | 2.25 | $2.47 \times 10^{-2}$ |
| poly(Segment, 9)9 × Impulse(CoP-$H_{fGn}$*) × Response(CoP-$SD$) | $2.72 \times 10^2$ | $9.86 \times 10^1$ | 2.76 | $5.83 \times 10^{-3}$ |
| Impulse(CoP-$t_{MF}$) | $9.75 \times 10^{-1}$ | $3.84 \times 10^{-1}$ | 2.54 | $1.12 \times 10^{-2}$ |
| poly(Segment, 9)1 × Impulse(CoP-$t_{MF}$) | $1.73 \times 10^2$ | $8.05 \times 10^2$ | 2.15 | $3.17 \times 10^{-2}$ |
| poly(Segment, 9)2 × Impulse(CoP-$t_{MF}$) | $2.03 \times 10^2$ | $8.05 \times 10^2$ | 2.52 | $1.19 \times 10^{-2}$ |
| poly(Segment, 9)3 × Impulse(CoP-$t_{MF}$) | $4.14 \times 10^2$ | $8.05 \times 10^2$ | 5.14 | $2.71 \times 10^{-7}$ |
| poly(Segment, 9)4 × Impulse(CoP-$t_{MF}$) | $2.38 \times 10^2$ | $8.05 \times 10^2$ | 2.95 | $3.17 \times 10^{-3}$ |
| poly(Segment, 9)5 × Impulse(CoP-$t_{MF}$) | $4.73 \times 10^2$ | $8.05 \times 10^2$ | 5.87 | $4.41 \times 10^{-9}$ |
| poly(Segment, 9)6 × Impulse(CoP-$t_{MF}$) | $2.64 \times 10^2$ | $8.05 \times 10^2$ | 3.28 | $1.05 \times 10^{-3}$ |
| poly(Segment, 9)7 × Impulse(CoP-$t_{MF}$) | $4.91 \times 10^2$ | $8.05 \times 10^2$ | 6.10 | $1.08 \times 10^{-9}$ |
| poly(Segment, 9)8 × Impulse(CoP-$t_{MF}$) | $2.53 \times 10^2$ | $8.05 \times 10^2$ | 3.14 | $1.68 \times 10^{-3}$ |
| poly(Segment, 9)9 × Impulse(CoP-$t_{MF}$) | $5.72 \times 10^2$ | $8.05 \times 10^2$ | 7.11 | $1.21 \times 10^{-12}$ |
| Impulse(CoP-$t_{MF}$) × Response(CoP-$H_{fGn}$) | $-9.74 \times 10^{-1}$ | $4.52 \times 10^{-1}$ | -2.15 | $3.11 \times 10^{-2}$ |
| poly(Segment, 9)2 × Impulse(CoP-$t_{MF}$) × Response(CoP-$H_{fGn}$) | $-2.02 \times 10^2$ | $9.86 \times 10^1$ | -2.05 | $4.01 \times 10^{-2}$ |
| poly(Segment, 9)3 × Impulse(CoP-$t_{MF}$) × Response(CoP-$H_{fGn}$) | $-4.14 \times 10^2$ | $9.86 \times 10^1$ | -4.20 | $2.70 \times 10^{-5}$ |
| poly(Segment, 9)4 × Impulse(CoP-$t_{MF}$) × Response(CoP-$H_{fGn}$) | $-2.37 \times 10^2$ | $9.86 \times 10^1$ | -2.41 | $1.61 \times 10^{-2}$ |
| poly(Segment, 9)5 × Impulse(CoP-$t_{MF}$) × Response(CoP-$H_{fGn}$) | $-4.72 \times 10^2$ | $9.86 \times 10^1$ | -4.79 | $1.68 \times 10^{-6}$ |
| poly(Segment, 9)6 × Impulse(CoP-$t_{MF}$) × Response(CoP-$H_{fGn}$) | $-2.64 \times 10^2$ | $9.86 \times 10^1$ | -2.67 | $7.51 \times 10^{-3}$ |
| poly(Segment, 9)7 × Impulse(CoP-$t_{MF}$) × Response(CoP-$H_{fGn}$) | $-4.91 \times 10^2$ | $9.86 \times 10^1$ | -4.98 | $6.51 \times 10^{-7}$ |
| poly(Segment, 9)8 × Impulse(CoP-$t_{MF}$) × Response(CoP-$H_{fGn}$) | $-2.53 \times 10^2$ | $9.86 \times 10^1$ | -2.56 | $1.04 \times 10^{-2}$ |
| poly(Segment, 9)9 × Impulse(CoP-$t_{MF}$) × Response(CoP-$H_{fGn}$) | $-5.72 \times 10^2$ | $9.86 \times 10^1$ | -5.80 | $6.78 \times 10^{-9}$ |
| Response(CoP-$SD$*) | $-9.78 \times 10^{-1}$ | $3.63 \times 10^{-1}$ | -2.70 | $7.00 \times 10^{-3}$ |
| poly(Segment, 9)1 × Response(CoP-$SD$*) | $-1.73 \times 10^2$ | $7.87 \times 10^1$ | -2.20 | $2.78 \times 10^{-2}$ |
| poly(Segment, 9)2 × Response(CoP-$SD$*) | $-2.02 \times 10^2$ | $7.87 \times 10^1$ | -2.58 | $9.93 \times 10^{-3}$ |
| poly(Segment, 9)3 × Response(CoP-$SD$*) | $-4.14 \times 10^2$ | $7.87 \times 10^1$ | -5.27 | $1.38 \times 10^{-7}$ |
| poly(Segment, 9)4 × Response(CoP-SD*) | -2.37×10<sup>2</sup> | 7.87×10<sup>1</sup> | -3.02 | **2.54×10<sup>-3</sup>** |
| poly(Segment, 9)5 × Response(CoP-SD*) | -4.73×10<sup>2</sup> | 7.87×10<sup>1</sup> | -6.01 | **1.83×10<sup>-9</sup>** |
| poly(Segment, 9)6 × Response(CoP-SD*) | -2.64×10<sup>2</sup> | 7.87×10<sup>1</sup> | -3.35 | **7.97×10<sup>-4</sup>** |
| poly(Segment, 9)7 × Response(CoP-SD*) | -4.91×10<sup>2</sup> | 7.87×10<sup>1</sup> | -6.25 | **4.20×10<sup>-10</sup>** |
| poly(Segment, 9)8 × Response(CoP-SD*) | -2.53×10<sup>2</sup> | 7.87×10<sup>1</sup> | -3.22 | **1.30×10<sup>-3</sup>** |
| poly(Segment, 9)9 × Response(CoP-SD*) | -5.73×10<sup>2</sup> | 7.87×10<sup>1</sup> | -7.28 | **3.39×10<sup>-13</sup>** |
| Response(CoP-t<sub>MF</sub>) | -5.57×10<sup>-1</sup> | 2.66×10<sup>-1</sup> | -2.09 | **3.64×10<sup>-2</sup>** |
| poly(Segment, 9)9 × Response(CoP-t<sub>MF</sub>) | 1.15×10<sup>2</sup> | 5.69×10<sup>1</sup> | 2.02 | **4.33×10<sup>-2</sup>** |

<sup>a</sup>Fitted model: IRF ~ (EC + CVD + DF\_CVD)\*poly(segment,9)\*Impulse\*Response + (1|Participant)
<sup>b</sup>Boldfaced values indicate statistical significance at the the alpha level of 0.05.

**Table 6.** Significant effects of Distance from Comfortable Viewing Distance (CVD) from mixed-effect regression modeling^a^ of impulse-response functions incorporating CoP-*t*_MF_ along with the prior set of CoM-*H*_fGn_, CoP-*SD*, and CoP-*H*_fGn_ (see Section 3.2.4.). ‘*’ after *H*_fGn_ and *SD* indicate the impulse-response functions from the second vector autoregression.

| Effect | *B* | *SE* | *t* | *P*<sup>b</sup> |
| --- | --- | --- | --- | --- |
| CVD × poly(Segment, 9)9 × Impulse(CoP-$H_{fGn}$*) | $2.32 \times 10^2$ | $1.15 \times 10^2$ | 2.02 | **$4.30 \times 10^{-1}$** |
| CVD × poly(Segment, 9)3 × Impulse(CoP-$t_{MF}$) | $-3.53 \times 10^2$ | $1.15 \times 10^2$ | -3.07 | **$2.12 \times 10^{-2}$** |
| CVD × poly(Segment, 9)4 × Impulse(CoP-$t_{MF}$) | $-2.47 \times 10^2$ | $1.15 \times 10^2$ | -2.15 | **$3.17 \times 10^{-1}$** |
| CVD × poly(Segment, 9)5 × Impulse(CoP-$t_{MF}$) | $-3.70 \times 10^2$ | $1.15 \times 10^2$ | -3.23 | **$1.26 \times 10^{-2}$** |
| CVD × poly(Segment, 9)6 × Impulse(CoP-$t_{MF}$) | $-2.63 \times 10^2$ | $1.15 \times 10^2$ | -2.29 | **$2.22 \times 10^{-1}$** |
| CVD × poly(Segment, 9)7 × Impulse(CoP-$t_{MF}$) | $-4.10 \times 10^2$ | $1.15 \times 10^2$ | -3.57 | **$3.52 \times 10^{-4}$** |
| CVD × poly(Segment, 9)8 × Impulse(CoP-$t_{MF}$) | $-2.33 \times 10^2$ | $1.15 \times 10^2$ | -2.03 | **$4.27 \times 10^{-2}$** |
| CVD × poly(Segment, 9)9 × Impulse(CoP-$t_{MF}$) | $-4.93 \times 10^2$ | $1.15 \times 10^2$ | -4.30 | **$1.74 \times 10^{-5}$** |
| CVD × poly(Segment, 9)3 × Impulse(CoP-$t_{MF}$) × Response(CoP-$H_{fGn}$) | $3.53 \times 10^2$ | $1.41 \times 10^2$ | 2.51 | **$1.21 \times 10^{-2}$** |
| CVD × poly(Segment, 9)5 × Impulse(CoP-$t_{MF}$) × Response(CoP-$H_{fGn}$) | $3.70 \times 10^2$ | $1.41 \times 10^2$ | 2.63 | **$8.50 \times 10^{-3}$** |
| CVD × poly(Segment, 9)7 × Impulse(CoP-$t_{MF}$) × Response(CoP-$H_{fGn}$) | $4.10 \times 10^2$ | $1.41 \times 10^2$ | 2.92 | **$3.55 \times 10^{-3}$** |
| CVD × poly(Segment, 9)9 × Impulse(CoP-$t_{MF}$) × Response(CoP-$H_{fGn}$) | $4.93 \times 10^2$ | $1.41 \times 10^2$ | 3.50 | **$4.58 \times 10^{-4}$** |
| CVD × poly(Segment, 9)3 × Response(CoP-$SD$*) | $3.53 \times 10^2$ | $1.10 \times 10^2$ | 3.22 | **$1.29 \times 10^{-3}$** |
| CVD × poly(Segment, 9)4 × Response(CoP-$SD$*) | $2.47 \times 10^2$ | $1.10 \times 10^2$ | 2.25 | **$2.44 \times 10^{-2}$** |
| CVD × poly(Segment, 9)5 × Response(CoP-$SD$*) | $3.70 \times 10^2$ | $1.10 \times 10^2$ | 3.38 | **$7.34 \times 10^{-4}$** |
| CVD × poly(Segment, 9)6 × Response(CoP-$SD$*) | $2.63 \times 10^2$ | $1.10 \times 10^2$ | 2.40 | **$1.65 \times 10^{-2}$** |
| CVD × poly(Segment, 9)7 × Response(CoP-$SD$*) | $4.10 \times 10^2$ | $1.10 \times 10^2$ | 3.74 | **$1.84 \times 10^{-4}$** |
| CVD × poly(Segment, 9)8 × Response(CoP-$SD$*) | $2.33 \times 10^2$ | $1.10 \times 10^2$ | 2.12 | **$3.37 \times 10^{-2}$** |
| CVD × poly(Segment, 9)9 × Response(CoP-$SD$*) | $4.93 \times 10^2$ | $1.10 \times 10^2$ | 4.50 | **$6.89 \times 10^{-6}$** |

<sup>a</sup>Fitted model: $IRF \sim (EC + CVD + DF\_CVD) * \text{poly}(\text{segment}, 9) * \text{Impulse} * \text{Response} + (1 | \text{Participant})$
<sup>b</sup>Boldfaced values indicate statistical significance at the the alpha level of 0.05.

As visual precision demands increased, increases in multifractal nonlinearity predicted progressively more stable reductions in subsequent *SD* (Fig. 9A). That is, compared to 50-cm viewing condition, subsequent changes in *SD* showed progressively less oscillation and more consistent reductions in subsequent *SD* with progressively more deviation from 50-cm viewing distance. This effect was indicated by the 9 ^th^-order polynomials reflecting the impulse from CoP-*t*_MF_ with increasing distance from 50 cm (DF_CVD × 9^th^-order Segment × Impulse(CoP-*t*_MF_), polynomial coefficients from linear to nonic, *P*s < 0.05, Table 7) and the response from CoP-*SD* with increasing distance from 50 cm (DF_CVD × 9^th^-order Segment × Response(CoP-*SD*\*), polynomial coefficients from linear to nonic, *P*s < 0.05, Table 7).

**Table 7.** Significant effects of Eyes Closed (EC) from mixed-effect regression modeling^a^ of impulse-response functions incorporating CoP-*t*_MF_ along with the prior set of CoM-*H*_fGn_, CoP-*SD*, and CoP-*H*_fGn_ (see Section 3.2.2.). ‘*’ after *H*_fGn_ and *SD* indicate the impulse-response functions from the second vector autoregression.

| Effect | *B* | *SE* | *t* | *P*<sup>b</sup> |
| --- | --- | --- | --- | --- |
| DF_CVD × Impulse(CoP-$H_{fGn}$*) | $7.10 \times 10^{-3}$ | $2.29 \times 10^{-3}$ | 3.09 | **$1.99 \times 10^{-3}$** |
| DF_CVD × poly(Segment, 9)5 × Impulse(CoP-$H_{fGn}$*) | $1.18 \times 10^0$ | $5.06 \times 10^{-1}$ | 2.33 | **$1.99 \times 10^{-2}$** |
| DF_CVD × poly(Segment, 9)7 × Impulse(CoP-$H_{fGn}$*) | $1.12 \times 10^0$ | $5.06 \times 10^{-1}$ | 2.22 | **$2.63 \times 10^{-2}$** |
| DF_CVD × poly(Segment, 9)9 × Impulse(CoP-$H_{fGn}$*) | $1.40 \times 10^0$ | $5.06 \times 10^{-1}$ | 2.77 | **$5.63 \times 10^{-3}$** |
| DF_CVD × Impulse(CoP-$H_{fGn}$*) × Response(CoP-$H_{fGn}$) | $-7.20 \times 10^{-3}$ | $2.81 \times 10^{-3}$ | -2.56 | **$1.04 \times 10^{-2}$** |
| DF_CVD × poly(Segment, 9)9 × Impulse(CoP-$H_{fGn}$*) × Response(CoP-$H_{fGn}$) | $-1.38 \times 10^0$ | $6.19 \times 10^{-1}$ | -2.23 | **$2.60 \times 10^{-2}$** |
| DF_CVD × Impulse(CoP-$H_{fGn}$*) × Response(CoP-$SD$) | $-7.09 \times 10^{-3}$ | $2.81 \times 10^{-3}$ | -2.52 | **$1.16 \times 10^{-2}$** |
| DF_CVD × poly(Segment, 9)9 × Impulse(CoP-$H_{fGn}$*) × Response(CoP-$SD$) | $-1.40 \times 10^0$ | $6.19 \times 10^{-1}$ | -2.26 | **$2.38 \times 10^{-2}$** |
| DF_CVD × poly(Segment, 9)1 × Impulse(CoP-$t_{MF}$) | $-1.01 \times 10^0$ | $5.06 \times 10^{-1}$ | -2.00 | **$4.60 \times 10^{-2}$** |
| DF_CVD × poly(Segment, 9)2 × Impulse(CoP-$t_{MF}$) | $-1.20 \times 10^0$ | $5.06 \times 10^{-1}$ | -2.37 | **$1.78 \times 10^{-2}$** |
| DF_CVD × poly(Segment, 9)3 × Impulse(CoP-$t_{MF}$) | $-1.68 \times 10^0$ | $5.06 \times 10^{-1}$ | -3.33 | **$8.64 \times 10^{-4}$** |
| DF_CVD × poly(Segment, 9)4 × Impulse(CoP-$t_{MF}$) | $-1.51 \times 10^0$ | $5.06 \times 10^{-1}$ | -2.99 | **$2.80 \times 10^{-3}$** |
| DF_CVD × poly(Segment, 9)5 × Impulse(CoP-$t_{MF}$) | $-2.03 \times 10^0$ | $5.06 \times 10^{-1}$ | -4.02 | **$5.79 \times 10^{-5}$** |
| DF_CVD × poly(Segment, 9)6 × Impulse(CoP-$t_{MF}$) | $-1.57 \times 10^0$ | $5.06 \times 10^{-1}$ | -3.11 | **$1.90 \times 10^{-3}$** |
| DF_CVD × poly(Segment, 9)7 × Impulse(CoP-$t_{MF}$) | $-2.19 \times 10^0$ | $5.06 \times 10^{-1}$ | -4.33 | **$1.49 \times 10^{-5}$** |
| DF_CVD × poly(Segment, 9)8 × Impulse(CoP-$t_{MF}$) | $-1.39 \times 10^0$ | $5.06 \times 10^{-1}$ | -2.75 | **$5.87 \times 10^{-3}$** |
| DF_CVD × poly(Segment, 9)9 × Impulse(CoP-$t_{MF}$) | $-2.64 \times 10^0$ | $5.06 \times 10^{-1}$ | -5.22 | **$1.84 \times 10^{-7}$** |
| DF_CVD × poly(Segment, 9)3 × Impulse(CoP-$t_{MF}$) × Response(CoP-$H_{fGn}$) | $1.68 \times 10^0$ | $6.19 \times 10^{-1}$ | 2.72 | **$6.56 \times 10^{-3}$** |
| DF_CVD × poly(Segment, 9)4 × Impulse(CoP-$t_{MF}$) × Response(CoP-$H_{fGn}$) | $1.51 \times 10^0$ | $6.19 \times 10^{-1}$ | 2.44 | **$1.47 \times 10^{-2}$** |
| DF_CVD × poly(Segment, 9)5 × Impulse(CoP-$t_{MF}$) × Response(CoP-$H_{fGn}$) | $2.03 \times 10^0$ | $6.19 \times 10^{-1}$ | 3.28 | **$1.03 \times 10^{-3}$** |
| DF_CVD × poly(Segment, 9)6 × Impulse(CoP-$t_{MF}$) × Response(CoP-$H_{fGn}$) | $1.57 \times 10^0$ | $6.19 \times 10^{-1}$ | 2.53 | **$1.13 \times 10^{-2}$** |
| DF_CVD × poly(Segment, 9)7 × Impulse(CoP-$t_{MF}$) × Response(CoP-$H_{fGn}$) | $2.19 \times 10^0$ | $6.19 \times 10^{-1}$ | 3.53 | **$4.10 \times 10^{-4}$** |
| DF_CVD × poly(Segment, 9)8 × Impulse(CoP-$t_{MF}$) × Response(CoP-$H_{fGn}$) | $1.39 \times 10^0$ | $6.19 \times 10^{-1}$ | 2.25 | **$2.46 \times 10^{-2}$** |
| DF_CVD × poly(Segment, 9)9 × Impulse(CoP-$t_{MF}$) × Response(CoP-$H_{fGn}$) | $2.64 \times 10^0$ | $6.19 \times 10^{-1}$ | 4.26 | **$2.09 \times 10^{-5}$** |
| DF_CVD × poly(Segment, 9)1 × Response(CoP-$SD$*) | $1.01 \times 10^0$ | $4.94 \times 10^{-1}$ | 2.04 | **$4.13 \times 10^{-2}$** |
| DF_CVD × poly(Segment, 9)2 × Response(CoP-$SD$*) | $1.20 \times 10^0$ | $4.94 \times 10^{-1}$ | 2.43 | **$1.53 \times 10^{-2}$** |
| DF_CVD × poly(Segment, 9)3 × Response(CoP-$SD$*) | $1.68 \times 10^0$ | $4.94 \times 10^{-1}$ | 3.41 | **$6.56 \times 10^{-4}$** |
| DF_CVD × poly(Segment, 9)4 × Response(CoP-$SD$*) | $1.51 \times 10^0$ | $4.94 \times 10^{-1}$ | 3.06 | **$2.22 \times 10^{-3}$** |
| DF_CVD × poly(Segment, 9)5 × Response(CoP-$SD$*) | $2.03 \times 10^0$ | $4.94 \times 10^{-1}$ | 4.11 | **$3.90 \times 10^{-5}$** |
| DF_CVD × poly(Segment, 9)6 × Response(CoP-$SD$*) | $1.57 \times 10^0$ | $4.94 \times 10^{-1}$ | 3.18 | **$1.48 \times 10^{-3}$** |
| DF_CVD × poly(Segment, 9)7 × Response(CoP-$SD$*) | $2.19 \times 10^0$ | $4.94 \times 10^{-1}$ | 4.43 | **$9.42 \times 10^{-6}$** |
| DF_CVD × poly(Segment, 9)8 × Response(CoP-$SD$*) | $1.39 \times 10^0$ | $4.94 \times 10^{-1}$ | 2.82 | **$4.82 \times 10^{-3}$** |
| DF_CVD × poly(Segment, 9)9 × Response(CoP-$SD$*) | $2.64 \times 10^0$ | $4.94 \times 10^{-1}$ | 5.34 | **$9.52 \times 10^{-8}$** |
| DF_CVD × Response(CoP-$t_{MF}$) | $-7.30 \times 10^{-3}$ | $1.62 \times 10^{-3}$ | -4.50 | **$6.82 \times 10^{-6}$** |

<sup>a</sup>Fitted model: $IRF \sim (EC + CVD + DF\_CVD) * \text{poly}(\text{segment}, 9) * \text{Impulse} * \text{Response} + (1 | \text{Participant})$
<sup>b</sup>Boldfaced values indicate statistical significance at the the alpha level of 0.05.

#### 3.2.2. Closing eyes led multifractal nonlinearity to predict subsequent increases in SD of postural fluctuations (Hypothesis-5b)

Compared to the eyes-open conditions, the closed-eyes condition led multifractal nonlinear to predict more increases in *SD* of postural fluctuations in 3 to 7 segments ahead and reductions in *SD* in 8 to 10 segments ahead (Fig. 9A). However, this late reduction in subsequent *SD* was consistently weaker than the reduction of *SD* in the eyes-open conditions. This effect was indicated by the 9^th^-order polynomials reflecting the impulse from CoP-*t*_MF_ with eyes closed (EC × Impulse(CoP-*t*_MF_), *B* = 2.26×10^0^, *P* = 1.31×10^-4^; EC × 9^th^-order Segment × Impulse(CoP-*t*_MF_), polynomial coefficients from linear to nonic, *P*s < 0.05, Table 8) and the response from CoP-*SD* with eyes closed (EC × Response(CoP-*SD*\*), *B* = −2.26×10^0^, *P* = 1.86×10 ^–4^; EC × 9^th^-order Segment × Response(CoP-*SD*\*), polynomial coefficients from linear to nonic except quadratic and octic, *P*s < 0.05).

**Table 8.** Significant effects of Distance from Comfortable Viewing Distance (DF_CVD) from mixed-effect regression modeling^a^ of impulse-response functions incorporating CoP-*t*_MF_ along with the prior set of CoM-*H*_fGn_, CoP-*SD*, and CoP-*H*_fGn_ (see Section 3.2.3.). ‘*’ after *H*_fGn_ and *SD* indicate the impulse-response functions from the second vector autoregression.

| Effect | *B* | *SE* | *t* | *P*<sup>b</sup> |
| --- | --- | --- | --- | --- |
| EC × Impulse(CoP-$H_{fGn}$*) | $1.75 \times 10^0$ | $5.91 \times 10^{-1}$ | 2.95 | **$3.15 \times 10^{-3}$** |
| EC × poly(Segment, 9)9 × Impulse(CoP-$H_{fGn}$*) | $2.84 \times 10^2$ | $1.30 \times 10^2$ | 2.18 | **$2.92 \times 10^{-2}$** |
| EC × Impulse(CoP-$H_{fGn}$*) × Response(CoP-$H_{fGn}$) | $-1.67 \times 10^0$ | $7.24 \times 10^{-1}$ | -2.31 | **$2.11 \times 10^{-2}$** |
| EC × Impulse(CoP-$H_{fGn}$*) × Response(CoP-$SD$) | $-1.75 \times 10^0$ | $7.24 \times 10^{-1}$ | -2.41 | **$1.59 \times 10^{-2}$** |
| EC × Impulse(CoP-$t_{MF}$) | $2.26 \times 10^0$ | $5.91 \times 10^{-1}$ | 3.83 | **$1.31 \times 10^{-4}$** |
| EC × poly(Segment, 9)1 × Impulse(CoP-$t_{MF}$) | $-3.52 \times 10^2$ | $1.30 \times 10^2$ | -2.70 | **$7.00 \times 10^{-3}$** |
| EC × poly(Segment, 9)2 × Impulse(CoP-$t_{MF}$) | $-2.59 \times 10^2$ | $1.30 \times 10^2$ | -1.98 | **$4.74 \times 10^{-2}$** |
| EC × poly(Segment, 9)3 × Impulse(CoP-$t_{MF}$) | $-2.82 \times 10^2$ | $1.30 \times 10^2$ | -2.17 | **$3.03 \times 10^{-2}$** |
| EC × poly(Segment, 9)4 × Impulse(CoP-$t_{MF}$) | $-3.84 \times 10^2$ | $1.30 \times 10^2$ | -2.94 | **$3.24 \times 10^{-3}$** |
| EC × poly(Segment, 9)5 × Impulse(CoP-$t_{MF}$) | $-3.39 \times 10^2$ | $1.30 \times 10^2$ | -2.60 | **$9.34 \times 10^{-3}$** |
| EC × poly(Segment, 9)6 × Impulse(CoP-$t_{MF}$) | $-3.59 \times 10^2$ | $1.30 \times 10^2$ | -2.75 | **$5.92 \times 10^{-3}$** |
| EC × poly(Segment, 9)7 × Impulse(CoP-$t_{MF}$) | $-4.44 \times 10^2$ | $1.30 \times 10^2$ | -3.41 | **$6.58 \times 10^{-4}$** |
| EC × poly(Segment, 9)8 × Impulse(CoP-$t_{MF}$) | $-2.59 \times 10^2$ | $1.30 \times 10^2$ | -1.99 | **$4.71 \times 10^{-2}$** |
| EC × poly(Segment, 9)9 × Impulse(CoP-$t_{MF}$) | $-5.62 \times 10^2$ | $1.30 \times 10^2$ | -4.31 | **$1.62 \times 10^{-5}$** |
| EC × Impulse(CoP-$t_{MF}$) × Response(CoP-$t_{MF}$) | $-2.26 \times 10^0$ | $7.24 \times 10^{-1}$ | -3.12 | **$1.80 \times 10^{-3}$** |
| EC × poly(Segment, 9)1 × Impulse(CoP-$t_{MF}$) × Response(CoP-$t_{MF}$) | $3.51 \times 10^2$ | $1.60 \times 10^2$ | 2.20 | **$2.78 \times 10^{-2}$** |
| EC × poly(Segment, 9)4 × Impulse(CoP-$t_{MF}$) × Response(CoP-$H_{fGn}$) | $3.83 \times 10^2$ | $1.60 \times 10^2$ | 2.40 | **$1.63 \times 10^{-2}$** |
| EC × poly(Segment, 9)5 × Impulse(CoP-$t_{MF}$) × Response(CoP-$H_{fGn}$) | $3.39 \times 10^2$ | $1.60 \times 10^2$ | 1.12 | **$3.39 \times 10^{-2}$** |
| EC × poly(Segment, 9)6 × Impulse(CoP-$t_{MF}$) × Response(CoP-$H_{fGn}$) | $3.59 \times 10^2$ | $1.60 \times 10^2$ | 2.25 | **$2.48 \times 10^{-2}$** |
| EC × poly(Segment, 9)7 × Impulse(CoP-$t_{MF}$) × Response(CoP-$H_{fGn}$) | $4.44 \times 10^2$ | $1.60 \times 10^2$ | 2.78 | **$5.44 \times 10^{-3}$** |
| EC × poly(Segment, 9)9 × Impulse(CoP-$t_{MF}$) × Response(CoP-$H_{fGn}$) | $5.62 \times 10^2$ | $1.60 \times 10^2$ | 3.52 | **$4.35 \times 10^{-4}$** |
| EC × Response(CoP-$SD$*) | $-2.26 \times 10^0$ | $6.04 \times 10^{-1}$ | -3.74 | **$1.86 \times 10^{-4}$** |
| EC × poly(Segment, 9)1 × Response(CoP-$SD$*) | $3.51 \times 10^2$ | $1.33 \times 10^2$ | 2.64 | **$8.35 \times 10^{-3}$** |
| EC × poly(Segment, 9)3 × Response(CoP-$SD$*) | $2.83 \times 10^2$ | $1.33 \times 10^2$ | 2.12 | **$3.38 \times 10^{-2}$** |
| EC × poly(Segment, 9)4 × Response(CoP-$SD$*) | $3.84 \times 10^2$ | $1.33 \times 10^2$ | 2.88 | **$3.96 \times 10^{-3}$** |
| EC × poly(Segment, 9)5 × Response(CoP-$SD$*) | $3.39 \times 10^2$ | $1.33 \times 10^2$ | 2.54 | **$1.10 \times 10^{-2}$** |
| EC × poly(Segment, 9)6 × Response(CoP-$SD$*) | $3.59 \times 10^2$ | $1.33 \times 10^2$ | 2.69 | **$7.07 \times 10^{-3}$** |
| EC × poly(Segment, 9)7 × Response(CoP-$SD$*) | $4.44 \times 10^2$ | $1.33 \times 10^2$ | 3.33 | **$8.54 \times 10^{-4}$** |
| EC × poly(Segment, 9)9 × Response(CoP-$SD$*) | $5.63 \times 10^2$ | $1.33 \times 10^2$ | 4.22 | **$2.43 \times 10^{-5}$** |
| EC × Response(CoP-$t_{MF}$) | $-8.56 \times 10^{-1}$ | $4.18 \times 10^{-1}$ | -5.12 | **$3.13 \times 10^{-7}$** |

<sup>a</sup>Fitted model: $IRF \sim (EC + CVD + DF\_CVD) * poly(segment, 9) * Impulse * Response + (1 | Participant)$
<sup>b</sup>Boldfaced values indicate statistical significance at the the alpha level of 0.05.

#### 3.2.3. Increases in multifractal nonlinearity predicted subsequent reductions in monofractal evidence in CoP fluctuations (Hypothesis-5c)

Increases in multifractal nonlinearity prompted an initial increase in monofractal evidence 1 Segment ahead for all the eyes-open conditions, after which the subsequent change went to practically zero at 2 Segments ahead (Fig. 9B). All the eyes-open conditions showed subsequent changes in monofractal evidence oscillating from more negative to more positive change by turn beginning with 3 Segments ahead. At the extremes of precision demand—both lowest (50-cm) and highest (220- and 305-cm)—the oscillations were actually weakest, with subsequent changes in monofractal evidence almost always positive. At the intermediary precision demands (25- and 135-cm), subsequent changes in monofractal evidence oscillated much more and with progressively larger amplitude with progressive segments, indicating negative changes 5, 7, and 9 segments ahead and indicating positive changes 4, 6, 8, and 10 segments ahead.

This effect for 50-cm viewing condition was indicated by 9^th^-order polynomials reflecting the impulses of CoP-*t*_MF_ and of CVD × Impulse(CoP-*t*_MF_) noted in Section 3.2.1 and the response from CoP-*H*_fGn_ to impulse from *t*_MF_ (Impulse(CoP-*t*_MF_) × Response(CoP-*H*_fGn_), *B* = −9.74×10^-1^, *P* = 3.11×10^-2^; 9^th^-order Segment × Impulse(CoP-*t*_MF_) × Response(CoP-*H*_fGn_), polynomial coefficients from quadratic to nonic, *P*s < 0.05, Table 5; CVD × 9^th^-order Segment × Impulse(CoP-*t*_MF_) × Response(CoP-*H*_fGn_), polynomial coefficients from quartic to nonic, *P*s < 0.05, Table 6). The effect for varying precision demands was indicated by the 9^th^-order polynomials reflecting the impulses of CoP-*t*_MF_ and of DF_CVD × Impulse(CoP-*t*_MF_) noted in Section 3.2.1. and the response from CoP-*H*_fGn_ with Impulse(CoP-*t*_MF_) for values of DF_CVD (DF_CVD × 9^th^-order Segment × Impulse(CoP-*t*_MF_) × Response(CoP-*H*_fGn_), polynomial coefficients from cubic to nonic, *P*s < 0.05, Table 7).

The eyes-closed condition diverged from model-predictions for the eyes-open conditions. It exhibited a large positive change in subsequent monofractal evidence 2 segments ahead, followed by smaller but relatively sustained and non-oscillatory changes in subsequent segments. This effect was indicated by the 9 ^th^-order polynomials reflecting the impulses from CoP-*t*_MF_ and EC × Impulse(CoP-*t*_MF_) noted in Section 3.2.2. and the response from CoP-*H*_fGn_ with impulse from CoP-*t*_MF_ (EC × Impulse(CoP-*t*_MF_) × Response(CoP-*H*_fGn_), B = – 2.26×10^0^, *P* = 1.80×10^-3^; EC × 9^th^-order Segment × Impulse(CoP-*t*_MF_) × Response(CoP-*H*_fGn_), polynomial coefficients from linear to nonic except quadratic, cubic, and octic, *P*s < 0.05, Table 8).

#### 3.2.4. Increases in monofractal evidence predicted subsequent increases in multifractal nonlinearity but less so for higher visual precision demands (Hypothesis-5d)

An increase in monofractal evidence predicted an increase in multifractal nonlinearity on the first segment ahead, resembling the pattern of effects in Section 3.2.3. on the first segment ahead. Hence, it appeared that increases in monofractal evidence and multifractal nonlinearity predicted subsequent increases in each other, but only in the short term. However, this resemblance to the effects of multifractal nonlinearity on monofractal evidence in Section 3.2.3 ended here.

Later than 1 segment ahead, monofractal evidence predicted subsequent increases in multifractal nonlinearity for all the eyes-open conditions until 5 segments ahead (Fig. 10A). From 5 segments ahead and onward, subsequent changes in monofractal nonlinearity began oscillating with progressively larger amplitude, reaching negative values 6 segments ahead in all conditions except 50-cm and 220-cm viewing conditions. The greatest subsequent changes appeared in 25- and 130-cm viewing conditions. Oscillations dwindled in 50-cm and 220-cm viewing conditions and then reversed in 305-cm viewing condition. In summary, monofractal evidence looked much like the inverse of results in Section 3.2.3. That is, when multifractal nonlinearity had predicted subsequent increases or reductions in monofractal evidence (i.e., in Section 3.2.3.), monofractal evidence predicted reductions or increases, respectively, in multifractal nonlinearity.

The foregoing effects were indicated by 9^th^-order polynomials reflecting the impulse from CoP-*H*_fGn_ for all values of DF_CVD (9^th^-order Segment × Impulse(CoP-*H*_fGn_*), quintic, septic, and nonic polynomial coefficients, Table 7) and the response from CoP-*t*_MF_ for all values of DF_CVD (Response(CoP-*t*_MF_), *B* = − 5.57×10^-1^, *P* = 3.64×10^-2^; 9^th^-order Segment × Response(CoP-*t*_MF_), nonic polynomial coefficient, *P* < 0.05, Table 5; DF_CVD × Response(CoP-*t*_MF_), *B* = *B* = −7.30×10^-3^, *P* = 6.82×10^-6^; DF_CVD × 9^th^-order Segment × Response(CoP-*t*_MF_), polynomial coefficients from linear to nonic, Table S2).

As in Section 3.2.3, 50-cm viewing condition yielded greater subsequent changes in multifractal nonlinearity comparable to that in 220-cm viewing condition (Fig. 10A). These effects were indicated by the 9^th^-order polynomials reflecting the impulse of CoP-H_fGn_ for values of CVD (CVD × 9^th^-order Segment × Impulse(CoP-*H*_fGn_*), nonic polynomial coefficient, Table 6) and reflecting Response(CoP-*t*_MF_) for values of CVD (CVD × 9^th^-order Segment × Response(CoP-*t*_MF_), polynomial coefficients from linear to nonic, Table S2).

The eyes-closed conditions showed the weakest relationship between monofractal evidence and subsequent changes in multifractal nonlinearity. Besides positive subsequent changes at 1 and 4 segments ahead and a negative subsequent change 3 segments ahead, subsequent changes in monofractal evidence was near zero. These effects were indicated by the 9^th^-order polynomials reflecting the impulse from CoP-H_fGn_ for values of EC (EC × Impulse(*H*_fGn_*), *B* = 1.75×10^0^, *P* = 3.15×10^-3^; EC × 9^th^-order Segment × Impulse(CoP-*H*_fGn_*), nonic polynomial coefficient, *P* < 0.05, Table 8) and reflecting the response from CoP-*t*_MF_ for values of EC (EC × Response(CoP-*t*_MF_), *B* = −8.56×10^-1^, *P* = 3.13×10^-7^; EC × 9^th^-order Segment × Response(CoP-*t*_MF_), polynomial coefficients from linear to nonic, Table S2).

#### 3.2.5. Increases in monofractal evidence predicted subsequent increases in SD for lower visual-precision demands but subsequent decreases for only the highest visual-precision demands (Hypothesis-5e)

Following an increase in monofractal evidence, the subsequent changes in *SD* were negative for all cases during the first 3 segments ahead—except the 25-cm viewing condition, which exhibited a subsequent increase in *SD* on the third segment ahead (Fig. 10B). Subsequently, the subsequent changes in SD oscillated for all the eyes-open conditions, remaining mostly negative for the greater precision demand (220- and 305-cm viewing conditions), but oscillating between positive and negative for intermediate precision demands (25- and 135-cm viewing conditions), with larger oscillations for the weaker precision demands (i.e., 25-cm). These effects were indicated by the 9^th^-order polynomials reflecting the response from CoP-*SD* for all values of DF_CVD (DF_CVD × Impulse(CoP-*H*_fGn_) × Response(CoP-*SD*), *B* = −7.09×10^-3^, *P* = 1.16×10^-2^; DF_CVD × 9^th^-order Segment × Impulse(CoP-*H*_fGn_) × Response(CoP-*SD*), nonic polynomial coefficient, *P* < 0.05, Table 7).

Meanwhile 50-cm viewing condition and eyes-closed conditions predicted subsequent increases in *SD* throughout beginning at 5 and 4 segments ahead, respectively (Fig. 10B). These effects were indicated by the 9^th^-order polynomials reflecting the response from CoP-*SD* for all values of CVD (CVD × 9^th^-order Segment × Impulse(CoP-*H*_fGn_) × Response(CoP-*SD*) polynomial coefficients from linear to nonic, Table S2) and for all values of EC (EC × Impulse(CoP-*H*_fGn_) × Response(CoP-*SD*), *B* = −1.75×10^0^, *P* < 1.59×10^-2^; EC × × Impulse(CoP-*H*_fGn_) × Response(CoP-*SD*), polynomial coefficients from linear to nonic, Table S2).

#### 3.2.6. Increases in SD predicted subsequent reductions in multifractal nonlinearity, with progressively larger reduction for progressively higher visual precision demands and with greatest reduction for the eyes-closed condition (Hypothesis-5f)

Greater *SD* in postural fluctuation predicted subsequent reductions in multifractal nonlinearity. This reduction was smallest for 50-cm viewing condition and progressively larger for conditions with higher visual precision demands (Fig. 11). These effects were carried entirely by linear components of the polynomial for the responses from CoP-*t*_MF_ for DF_CVD (Response(CoP-*t*_MF_), *B* = −5.57×10^-1^, *P* = 3.64×10^-2^; 9^th^-order Segment × Response(CoP-*t*_MF_), linear coefficient, *P* < 0.05, Table 5; DF_CVD × Response(CoP-*t*_MF_), *B* = −7.30×10^-3^, *P* = 6.82×10^-6^; DF_CVD × 9^th^-order Segment × Response(CoP-*t*_MF_), linear to nonic coefficients, Table S2) and CVD (CVD × Response(CoP-*t*_MF_), *B* = 1.7185×10^-2^, *P* = 2.14×10^1^; CVD × 9^th^-order Segment × Response(CoP-*t*_MF_), linear to nonic coefficients, Table S2).

The eyes-closed condition prompted the largest subsequent reductions in multifractal nonlinearity following increases in *SD* (EC × Response(CoP-*t*_MF_), *B* = −2.14×10^1^, *P* = 2.14×10^1^; EC × 9^th^-order Segment × Response(CoP-*t*_MF_), linear to nonic coefficients, Table S2, Fig. 11).

## 4. Discussion

We tested five hypothesis concerning how visual precision demands might moderate intrapostural interactivity. We predicted that: Standing quietly with the eyes closed would exhibit weaker intrapostural interactivity (Hypothesis-1). Monofractal evidence in CoM and CoP, CoM-*H*_fGn_ and CoP-*H*_fGn_, respectively, would self-correct over time (Hypothesis-2). An inverse relationship between monofractality and *SD* over time, that is, that increases in CoM-*H*_fGn_ and CoP-*H*_fGn_ would prompt subsequent reductions in CoP-*SD* (Hypothesis-3a) and that increases in CoP-*SD* would prompt subsequent reductions in CoM-*H*_fGn_ and CoP-*H*_fGn_ (Hypothesis-3b). These intrapostural interactions in the 50-cm viewing condition would most closely resemble intrapostural interactions in the eyes-closed condition (Hypothesis-4). Increases in multifractal nonlinearity CoP-*t*_MF_ would displace the role and response of monofractal evidence CoP-*H*_fGn_ appearing in Hypotheses-3a and −4, leaving monofractal evidence to show less precision-dependent reduction of CoP-*SD* and to contribute to multifractality for lower visual precision demands (Hypothesis-5). Results supported all hypotheses with the only exception being the failure of CoM-*H*_fGn_ to participate in the relationships predicted in Hypothesis-3. We discuss the implications of these findings below.

### 4.1. Ratcheting up our lens from monofractality to multifractality allows movement science to make contact with the nonlinear interactions across scales entailed by goal-directed and task-dependent behavior

This work vindicates the suggestion that multifractal dynamics might underlie the coordination supporting postural control. Multifractal dynamics and the mathematical, geometrical framework for investigating this proposal remains a novel and underdeveloped theme in movement science. Elements of this framework have been successfully introduced in terms of the popularization of monofractal geometry (e.g., 1/*f* noise as a key feature of movement variability; Newell & Jordan, 2007; Vaillancourt & Newell, 2002). However, the more recent calls have been to ratchet the modeling up to a new precision level. We briefly review the bigger picture below, on the way to explaining how this work supports a longer-range trajectory of developing multifractal perspectives on movement science. The outcome of these deliberations will be twofold. Monofractal modeling is an effective first step into multifractal modeling so far as multifractality follows from trial-by-trial variation in monofractal structure. However, monofractal modeling has limits, and the further modeling of multifractal spectra with the multifractal elaborations of monofractal modeling offers a deeper way to make inferential contact with nonlinear interactions across scales. The value of this work lies in showing that elaborating multifractal modeling beyond simply monofractal modeling—whether in isolation or sequence— reveals new insights into how postural control resolves the intermodality of organism-wide perception-action.

First, monofractal modeling has always been an excellent first step, but notably not the last step. Critically for motivating multifractal perspectives, the relationship between monofractal evidence and ‘nonlinearity’ is not as deep as initially hoped (Ihlen & Vereijken, 2010; Kelty-Stephen & Wallot, 2017). Yes, the power-law shape implicated in 1/f noise or any single fractal dimension is not linear, and yes, the model system once heralded as an explanation for 1/f noise (i.e., self-organized criticality) did enlist an interaction across behaviors situated on two different scales (Bak et al., 1987; Jensen, 1998). However, monofractal evidence has been insufficient on two fronts: it neither depends necessarily on self-organized criticality nor does it reach the full scope of academic aspirations motivating much of its interest. To the first point, the appearance of a single power law in our measurements is entirely consistent with and predictable from linear models that do not necessarily have any interactions across scales (Granger and Joyeux, 1980; Wagenmakers et al., 2004). To the second point, the theoretical discourse, particularly in ecological-psychological corners, has long sought to implicate multiple (i.e., more than two and even potentially ‘all’) scales of biological and task variability (Johnston & Turvey, 1980; Turvey & Fitzpatrick, 1993; Turvey, 2009; Wagman et al., 2016a, 2016b, 2017; Wagman & Miller, 2003; Wagman & Stoffregen, 2020). To sum up, self-organized criticality is just one case of a family of cascade models explicitly situating causal status in nonlinear interactions across scales, and choosing monofractal modeling to the exclusion of multifractal considerations risks missing that goal-directed behavior rests on cascade-driven causality beyond self-organized criticality (Stephen et al., 2012).

The issue of multifractal vs. monofractal modeling is about the lens we use to investigate goal-directed behavior. Monofractal modeling can provide evidence consistent with cascade-like nonlinear interactions across scales—cascades are one important source of monofractal structure (Mandelbrot, 1982). However, an isolated case of monofractal evidence is nothing that the linear model cannot predict (Granger and Joyeux, 1980), and any choice to interpret monofractal evidence as arising from cascades requires ruling out the null hypothesis that it arises from the linear model (Wagenmakers et al., 2004). The linear model’s capacity to predict monofractal evidence is difficult to imagine as a plausible alignment of cause. Specifically, it appeals to a model called ‘fractional integration’ contrived to permit all past timescales of behavior to contribute to the current behavior— with the puzzling entailment that none of these unbounded past behaviors interact with past behaviors at other scales. Hence, for some researchers, the implausibility of causal interpretations from this linear model originating from 1/*f* noise may be enough to warrant interpreting monofractal analysis as satisfactory evidence of cascades. This compromise exemplifies what Gilden (2009) warned of as ‘gross opportunism’ for the analytical sake of fitting the time-series data.

Monofractal analysis can presume interactions across scales, but multifractal analysis allows us to estimate the strength of these interactions and specific roles of nonlinearity in supporting them. The mere absurdity of linear explanation of monofractality does not somehow make monofractal evidence a better lens on nonlinearity. Estimating the time-variation of monofractal evidence is the first step towards multifractality and beyond the linear model of goal-directed behavior. If goal-directed behavior were strictly linear, then time-variation of monofractal evidence should be without structure and incapable of predicting perception-action outcomes (e.g., Section 1.3.2.1). However, as we highlighted in Section 1.3.2.2., analyses that are explicitly multifractal can estimate a ‘multifractal spectrum’ directly without the intermediary step of many sequential monofractal analyses. This spectrum width covaries with what you would find with sequential monofractal analysis, but whereas the spectrum width glosses over the actual sequences in this temporal variation, the multifractal spectrum offers the most compact method to estimate the nonlinear across-scales interactivity of a single measured time series. Sequential multifractal estimation of this spectrum, that is, for multiple time series in an experiment-wide sequence, thus reveals how nonlinear interactions across scales change alongside ongoing measures of goal-directed behavior.

Monofractal evidence and multifractal nonlinearity inform about ‘temporal structure’ and ‘nonlinear support for that structure,’ respectively. We can examine how the monofractal evidence and multifractal nonlinearity both change alongside more universally interesting outcome measures. Hence, the multifractal approach allows human movement science a wider set of features for articulating how nonlinear interactions across scales could matter for dexterous goal-directed behavior. Certainly, monofractal evidence involves a power-law relationship that unfolds across multiple scales, and the ‘self-similarity’ of this form is evocative of an interaction among the implicated scales. But multifractal-spectrum width compared to surrogate spectra provides the non-overlapping insight about whether the similar behavior at all scales reflect actual links between those scales.

The advantage of multifractal modeling is that it resolves heterogeneity and continuity in single framework. A major shortcoming of the monofractal modeling has been that a single estimate cannot necessarily specify all wider patterning of a distributed system (Likens et al., 2019, 2015). ‘Self-similarity’ is certainly a feature of power-law relationships supporting fractal and multifractal modeling, but we do not think it warrants a denial of heterogeneity. Nonlinear dynamics are fully compatible with a difference and even competition (e.g., Kelso, 1995). As Bernstein (1967) suggested, “movements are not chains of details but structures which are differentiated into details; they are structurally whole” (p. 69). The challenge is to maintain the possibility that heterogeneity does not violate the organism’s structural integrity (Stoffregen et al., 2017). Multifractal modeling affects this balance and may provide further details about the nesting of behaviors within the organism. For instance, multifractal nonlinear is a feature of physiological functioning (Ivanov et al., 2001), but manipulations of the nested structure in the task context (e.g., Wagman et al., 2016a, 2016b, 2017; Wagman & Stoffregen, 2020) may prompt organisms to embody the nesting relationships quite apart from what physiological needs demand. Multifractal analysis can reveal how nested structure in movement variability is sensitive to the nested structure of the task context (Eddy & Kelty-Stephen, 2015; Stephen & Dixon, 2011).

This work shows the progression from monofractal to multifractal modeling in practice with postural control as a model system. The findings can be summarized in the following way. Through the monofractal lens, monofractal evidence suggests that 1/*f*-like ‘temporal structure’ appears to reduce the *SD* of postural fluctuations, suggesting as well that this ‘temporal structure’ self-corrects, keeping itself around some happy medium of the temporal structure without excess in either direction. However, the multifractal lens shows us that 1/*f*-like temporal correlations are equally likely to increase or reduce the *SD* of postural fluctuations. The growth or decay of *SD* can be better explained in terms of prior changes in how organisms blend behaviors across different timescales. Any capacity of the organism to stabilize its monofractality inheres in its reservoir of nonlinear interactions across scales.

### 4.2. Glimpses of a possible postural control policy for visually guided quiet stance

The present results offer insights into a control policy for postural stability that balances CoP monofractality with excess CoP-*SD*. If left to *SD* alone, posture would lean towards higher variability without clear bound: any increase in *SD* would predict subsequent increases, and those subsequent increases would, in turn, predict further subsequent increases, and so on. The predicted subsequent reductions in monofractality would then only serve to promote greater *SD*. It is only the corrective aspect of monofractality that might allow posture to rein in the self-promoting and unbounded *SD*. For instance, any reductions in monofractality following increases in *SD* might trigger subsequent increases in monofractality that would induce a negative check on *SD*. Note that this causal interpretation aims only to offer a possible control policy that these results could reflect. A rigorous test of this causal interpretation warrants manipulations of *SD* and CoP monofractality through a balance board or vibrotactile stimulation.

The present results echoes the past findings that explicit feedback to participants completing a perceptuomotor task can weaken temporal correlations in movement variability (Kelty-Stephen & Dixon, 2014; Kuznetsov & Wallot, 2011; Stephen & Hajnal, 2011). In the task of counting seconds by tapping a finger (Kuznetsov and Wallot, 2011), the feedback provided with each tap allowed offsetting deviations, preventing errors from propagating from one tap to the next. At first glance, this finding seems at odds with the present finding that better performance—standing more quietly with less sway *SD*—would follow from and contribute to stronger and not weaker temporal correlations. However, it is possible that, in postural tasks, greater sway is endogenous, implicit feedback signaling the postural system that corrections have been appropriately implemented. In this way, if some proportion of *SD* reflects sway that triggers postural corrections (e.g., Hagio et al., 2018), then the present findings would align with the past findings of feedback decorrelating movement variability.

This control policy may resolve long-standing questions about how fluctuations support movement stability, explicitly speaking to the ‘loss of complexity’ hypothesis: monofractality in sway might reflect stability, suggestive of young, healthy, and typically developing physiology. This hypothesis has proven provocative but controversial. Indeed, clinical research has found that temporal correlations might wander within but also beyond the monofractal range. Additionally, the clinical implications of sway monofractality, with some results indicating stronger temporal correlations in sway for younger and healthier participants (Duarte and Sternad, 2008; Thurner et al., 2002) and other results indicating the opposite (Ko and Newell, 2016; Lipsitz, 2002; Thurner et al., 2002). Meanwhile, work applying ‘white-noise’—that is, temporally uncorrelated—mechanical vibration to the feet found that this, by definition, non-fractal and so non-complex signal stabilized sway (Priplata et al., 2002; Priplata et al., 2003). In finding that greater temporal correlations were associated with subsequent reductions in sway *SD*, the present results align poorly with the finding that uncorrelated stimulation reduced sway.

Results have curiously diverged within the same research paradigms in this vein, making matters seem even more paradoxical. A reanalysis of Priplata et al.’s (2003) data yielded two details (Kelty-Stephen and Dixon, 2013). Firstly, white-noise stimulation reduced temporal correlations in sway. So, in an unhealthy level of complexity, decorating overly correlated fluctuations may be a useful clinical strategy. Secondly and less straightforwardly, this reanalysis showed that white-noise stimulation elicited a stronger sway reduction in participants exhibiting stronger temporal correlations. In a sense, white-noise stimulation seems to wipe out its efficacy by counteracting the very conditions of endogenous postural fluctuations that give it a stabilizing effect, which seemed quite puzzling. This work solves some of this puzzle by finding predictive effects of fluctuation patterns as evident in the association among concurrent variables for the same postural measurement series. The significant contributions of examining prior effects and subsequent responses are twofold: first, monofractality self-corrects, and second, stronger monofractality reduced CoP-*SD*. Together, these two points provide a framework in which the present findings align neatly with research on white-noise stimulation stabilizing posture (Priplata et al., 2002; Priplata et al., 2003) and explain how fractal temporal correlations could be sometimes stabilizing and sometimes destabilizing.

Self-correction of fractal temporal correlations has been the key feature missing from the portrayal of postural stability until now. Lack of self-correction would mean that temporal correlations and *SD* might push each other to opposite extremes, subverting postural stability (Chen et al., 2008; Rajachandrakumar et al., 2018). Posture lacking sufficient variability would be unstable and overly temporally correlated; posture with too much variability would be unstable and have too little temporal correlations. Hence, understanding self-correction in temporal correlations might then be one of the key directions for future work. Intraindividual differences in temporal correlations are well-known (Collins and De Luca, 1995, 1993) and often replicated (Gilfriche et al., 2018). The novelty of this study lies in recognizing that weaker temporal correlations at longer timescales might hold only on average, consisting of the ebbing and flowing of stronger temporal correlations at shorter timescales. Across 10-s segments within a single 120-s trial, the vector autoregression found a sequence of fleeting (e.g., 10-s) bouts of postural sway predicting subsequent alternations between more or less temporal correlations.

The exact basis of this alternation of temporal correlations warrants further investigation. The question of whether these spatial constraints govern self-correction of temporal correlations would benefit from rigorous test in the ‘rambling-trembling’ framework that recognizes that the fixed reference point anchoring CoP within the base of support drifts slowly (Zatsiorsky & Duarte, 1999, 2000). Thus, it is important to model the rise and fall of temporal correlations as a function of the distance between the fixed reference point and the edges of the base of support (Kelty-Stephen et al., 2020). The space that ‘rambling’ leaves open for stable, ‘trembling’ may govern how quickly temporal correlations alternate. Such modeling could add to the ongoing elaborations of the rambling-trembling framework to include visual constraints (Ferronato and Barela, 2011; Yamagata et al., 2019).

In summary, the present results offer a glimpse of how current nonlinear dynamical models of self-correction (Boyd et al., 2017) may play out in biological goal-oriented behavior. Naturally, the correlational analysis results are prone to mischaracterizing the measured variables as actual causal variables. Fractal exponents and *SD* are measurements commonly thought to be essential state variables in postural control. Certainly, fractal exponents are not inherently physiological features but emergent properties of the organism’s interaction with the task. However, the actual postural control parameters may be no less emergent from task constraints—one of the rare agreements between current cognitivist theorizing about visual attention (Vecera et al., 2014) and long-standing views in ecological psychology (Bardy et al., 1999).

### 4.3. Postural control policy through the multifractal lens

#### 4.3.1. Multifractal nonlinearity drives the monofractal results and reveals novel distinction between eyes-closed and 50-cm viewing distance

Our inclusion of multifractal nonlinearity for modeling postural control was devastating to our findings related to monofractal evidence. No matter the validity of calling sequential monofractal evidence a case of ‘multifractality’ (Ihlen, 2012), the nonlinearity estimable by comparing multifractal spectra between original and surrogate series knocked through our monofractal modeling like a meteorite through the statistical roof. Indeed, from Hypothesis-4 to Hypothesis-5, the differences apparent in impulse-response functions were stark: not one of the impulse-response functions found for testing up to Hypothesis-4 survived with the inclusion of impulse-response functions with multifractality nonlinearity. This disparity indicates how much the landscape of modeled relationships could vary. So, although the emergent control may not have the same labels for its control parameters, similar observed relationships could explain a host of previous results, as discussed above. Indeed, multifractal nonlinearity appeared to step into relationships with *SD* that were most similar in form. It predicted subsequent reductions in *SD* with greater visual-precision demands, and it also reduced following increases in *SD*. So, through Hypothesis-4, the effects of monofractal evidence likely carried the deeper effects of yet-unestimated multifractal nonlinearity.

‘Self-correcting’ of monofractal evidence was not a feature of monofractality itself but instead reflected a close relationship between monofractality and multifractal nonlinearity. As we noted above in Section 1.3.2.2., monofractal structure and multifractal nonlinearity are mathematically distinguishable but closely related. The former indicates a correlation across time, and the latter indicates how diversity in this correlation. Now we see that monofractal evidence’s self-correction depends on an ongoing mutual push and pull between monofractality and multifractality. Because of the nonlinearity inherent in the movement system, changes in monofractal temporal correlations depend on and reshape the profile of interactions across scales. Nonlinearity supports changes in monofractal scaling, and changes in monofractal scaling could stir up new nonlinearity. This push and pull have two exciting features. First, it appears to be a case of mutual tempering, as the regression model’s cubic polynomial gives voice to the oscillatory pattern of impulse-response functions. Second, it diminishes with greater visual precision demands in a suprapostural fixation task. Whether this relationship holds in other task settings is unknown, but it highlights how monofractal evidence from multifractal nonlinearity could be separate variables reflecting distinct dimensions in a parameter space governing goal-directed behavior (Saltzman and Munhall, 1992).

The relationship between monofractality and multifractality is unequal, as multifractality appears to predict monofractality to a greater extent than the extent to which monofractality predicts multifractality. Furthermore, the multifractal model revealed more than the monofractal model about the research design supporting the present dataset. For instance, the monofractal model predicted average changes in subsequent *SD* only in viewing conditions other than the 50-cm viewing condition. Including multifractality reshaped the impulse-response functions to reveal positive effects of monofractal evidence on subsequent *SD* for both the 50cm viewing and eyes-closed conditions. Besides reshaping the apparent predictive role of monofractal evidence, multifractal nonlinearity also predicted a subsequent increase in *SD* and even estimated unique effects, situating the 50-cm viewing condition as an intermediary between the SD-increasing effect in the eyes-closed condition and *SD*-reductive effect in higher-precision conditions.

These results offer the latest caveat against simple answers to a long-running curiosity for simple resolution of “whether more (multi)fractality is better” in all cases. The capacity of multifractal nonlinearity to predict task-sensitive changes in opposite directions within the same research design is not new (Bell et al., 2019; Mangalam and Kelty-Stephen, 2020). Hence, we see that abruptly changing the support for posture, particularly in removing the task prompt for participants to reach out and invest exploratory behavior beyond the bounds of support, may make multifractal nonlinearity a liability. When the priority is to draw degrees of freedom inward (e.g., to stand as still as possible), then the variability entailed by multifractal nonlinearity becomes a source of greater spatial instability as indicated by *SD*. Certainly, the profiles of a subsequent change in *SD* show the 50-cm viewing condition in between the eyes-closed and higher-precision conditions. This point is reassuring in so far as the 50-cm viewing distance does represent a case where the visible layout beyond the eyelids is available but not demanding exploratory engagement in the way that other viewing distances do. Mere current availability of the visible layout may be enough to coax degrees of freedom ‘outward.’

Prior remarks are not a conclusion but a proposal to inquire into the behavioral value of multifractality. What counts as ‘inward’ and how to chart ‘outwardness’ becomes an immediate challenge to such reasoning, but it may be a way to begin organizing a set of findings of a deep set of observables. If multifractal nonlinearity is pervasive support for context- and task-sensitive goal-directed behavior, then we need an open mind to the sensitivity of multifractal nonlinearity to various task parameters over different time scales. For instance, consider a seemingly more formal organizing theme of ‘perturbation.’ Changes in the availability of visual information certainly could be construed as a perturbation, but then quick conclusions that, for example, “perturbations make multifractality worse for motor outcomes” will be too fragile to hold—on a very short, fleeting timescale, multifractal nonlinearity can support a constriction of SD in motor performance—both in past research and in the ‘2 segments later’ predicted value for the present case of our eyes-closed condition.

We expect that the unique rooting of multifractal nonlinearity in interactions across scales makes it critical for task sensitivity. The apparent links between multifractal nonlinearity and performance outcomes are thus likely to change with the task. Indeed, a challenge is that tasks are always deeper than catch-all factors like perturbation. These seemingly task-general terms like perturbation do not speak to the relevance of multiple scales in a seemingly straightforward and simple task. For instance, we recognize that the short timescale of segment and trials in the present reanalysis sets a yet mostly undetermined boundary on how far these exact findings generalize. Not only does multifractality opens up the possibility of variability with time, we know for sure that longer timescales can show different relations between multifractal profiles and behavioral or clinical outcomes (Bell et al., 2019; Ivanov et al., 2001).

Similarly, related to timescales, one can argue that these results do not generalize to situations in which participants are not instructed to minimize sway or fixate gaze. However, we already know from past studies that multifractal nonlinearity carries significant predictive weight as well when participants can comfortably move as they prefer in tasks that do not prompt such tight restraint of the degrees of freedom (Carver et al., 2017; Carver & Kelty-Stephen, 2017; Jacobson et al., 2020; Kelty-Stephen & Dixon, 2014; Stephen et al., 2012; Teng et al., 2016). The imprint of multifractality on goal-directed behavior may change its form depending on task constraints (e.g., our reported difference between eyes-closed and 50-cm viewing conditions), but the role of multifractal nonlinearity will not simply vanish when the participants break out of instructions to restrict movement.

Contrived as task settings may be, they still constitute a thicket of nesting relationships and relationships across scales (e.g., Wimsatt, 1994). Our concern is not naturalism (multifractality remains a useful predictor of goal-directed behavior in outdoor, nonlaboratory settings; Teng et al., 2016). Instead, we are concerned that scientists are still delving into this issue of many nested constraints in a task—well beyond the explicit instructions and simple presentations of stimuli. For all the power of controlling task constraints, this control comes with critical blind spots. Indeed, focusing on a specific task manipulation makes ignoring other factors almost inevitable. This concern is not new (e.g., Wachtel, 1973), but we suspect that, given its sensitivity to interactions across scales, multifractal nonlinearity would serve as an essential flashlight into the nestings that support observed behavior—potentially even into nesting relationships that were not intentionally built into task settings.

The organism participating in a task setting is no less a causal thicket, and a relatively new direction in multifractal modeling is to examine how multifractal fluctuations might flow through the body as different parts differently make contact with different aspects of globally available information. This approach suggests a more constructive question of “what sort of flow of multifractal fluctuations supports perception?” Through this lens, there is no real sense in which multifractality is abstractly good or abstractly bad. We already have evidence that the flow of multifractal fluctuations across the body predicts the accuracy of perceptual judgments (Mangalam et al., 2020a, 2020b). Of course, we look forward to the eventual question of which kinds of flow are better than others and for which tasks. Our point, for now, is that multifractality might sooner be a language in which a bodywide architecture like tensegrity may speak in ways we can listen to and learn from. There might yet be a reason to consider whether multifractal fluctuations provide a useful domain for synergies whose manifestation in abundant variability is a good and expectable ‘bliss’ (Latash, 2020, 2012).

For instance, multifractal nonlinearity vindicates the long-cherished idea of vision as a form of action, even in this sparse setting. It is worth noting that pre-multifractal modeling of this dataset construed ‘visual precision demand’ as being highest for 25-cm and 50-cm viewing conditions (Lee et al., 2019, p. 434). Multifractal modeling of this data from the perspective of cascading interactions across scales vindicates a wholly distinct form of visual precision demands, which is the departure of viewing distances in either direction from a resting point for lens accommodation roughly 50-70 cm before the participants’ eyes (Iwasaki and Tawara, 2002; Korge and Krueger, 1984; Leibowitz and Owens, 1978; Rempel et al., 2007; Sakamoto et al., 2012). Beyond the idea that vision and action are simply coexisting functionally related processes (Hayhoe, 2017), the multifractal nonlinearity of posture is thus laying bare the bodywide entailments of precision prompted from the intentional engagement of the lens to fixate at a distance (e.g., Gibson, 1988; Kim & Turvey, 1999). Certainly, ecological approaches respect that whole organisms are worth more than just the lens to muster in the service of visual perception (Gibson, 1966, 1979). Multifractal nonlinearity offers a quantitative grasp of how perception-action thrives on interactions between organism-scaled events with lens-scaled events. Indeed, multifractal nonlinearity may provide a substantial advance beyond the presumed rigor of explanatory appeals to strictly reductive treatments of physiology, giving force to progress in the directions suggested by ecological psychology.

The capacity of multifractal nonlinearity as a core feature of the movement system’s task-sensitivity raises an important possibility about which question is more apt. In the first place, authors must present novel descriptors to layout their plain meaning. So, we have real sympathy with the ongoing challenges in understanding ‘what multifractality means.’ But we also see a real benefit in not overpromising in the directions of how successful or accurate ‘more fractal’ or ‘more multifractal’ goal-directed behavior will be. It might be more constructive to discuss multifractal nonlinearity in conjunction with any other fractal-related descriptors as quantitative flashlights can bring relationships nested across scales. So long as these relationships might go unnoticed in an experimental design intended for other purposes, they reflect not just poetic acknowledgments of other factors not yet explored—but also theoretical questions currently dormant. So, multifractal nonlinearity may eventually provide a road map for deconstructing our ongoing experimental procedures and learning how to critically evaluate and then to design nesting relationships for newer research and applications.

#### 4.3.2. Implications for stimulation-based rehabilitation

One of the most accessible potential avenues for application of multifractality is simulation-based rehabilitation in which the so-called ‘noise’ is used to nourish the movement system with more of its healthy or optimal variability (e.g., Cavanaugh et al., 2017). The preceding in Section 4.2. speaks to a discourse on simulation-based rehabilitation that, to date, remains poised mainly at the monofractal lens (Ducharme & van Emmerik, 2018; Kelty-Stephen & Dixon, 2013; Priplata et al., 2002; Priplata et al., 2003; Rhea et al., 2014; Sotirakis et al., 2020; Vaz et al., 2020). It perches at the frontiers of movement science, and its promise lies in envisioning that multiple monofractal profiles at one time—even if it is just the contrast between two monofractal profiles: one of the organisms and the other of the stimulation. However, the caveat on the interpretation of correlational analyses remains in force, as a strictly monofractal discourse risks failing to acknowledge any of possible multifractality-based variables. This work asserts clearly that, if the movement system thrives on stochastic fluctuations, then we might need to prescribe variable stimulation with no less multifractal structure.

The restructuring of our impulse-response functions and the expansion of our nonlinear-dynamical space for modeling goal-directed behavior resonate with the current calls for new directions in simulation-based rehabilitation. Recent work has examined the role of the auditory stimulus with elaborations beyond fractal structure in supporting gait (Hove et al., 2012; Hunt et al., 2014; Rodger and Craig, 2016; Uchitomi et al., 2013). Despite never explicitly naming ‘multifractal’ structure as focus, these studies propose to build gait-scaffolding stimulation from the foundation of biological patterns (e.g., gait and music) that marry multi-scale aspect of fractal fluctuations with complex phase relationships amongst participating timescales. This blend of features guarantees multifractality in these structures (Muñoz-Diosdado, 2005; Telesca and Lovallo, 2011). Likewise, if we study the postural system to identify factors that can minimize sway, it becomes evident that the postural system itself exhibits a multifractal structure. Indeed, the stochastic resonance that the scientific community has been trying to build might be more accessible by using multifractal stimulation.

The multifractal reanalysis could inform considerations about how endogenous fluctuations inform tailoring stimulation for rehabilitation, potentially moving it past the frontiers of monofractal formalism. The limitation of fractal temporal correlations is that fractal models are all too symmetrical, and postural sway is known to be intermittent (Lee et al., 2018, 2019). Monofractal stimulation may be ill-suited to the multifractal landscape of postural control. Future work could benefit from considering what multifractal configurations of posture will be most receptive to stimulation, and multifractal geometry may offer the plainest framework for tuning stimuli to multifractal control of posture. In pointing to the importance of variation in fractal temporal correlations, this work is a step towards understanding the multisensory aspect of posture and providing multisensory support for its clinical support and rehabilitation.

To sum up, the present findings on quietly standing and fixating situates intermodal use of information on multifractal foundations. Visually fixating brings a prestressed quiet stance into one with informational coupling with the visual stimulus. Past work has repeatedly implicated multifractal fluctuations in the head and upper torso for using visual information to organize action (Bell et al., 2019; Carver et al., 2017; Eddy and Kelty-Stephen, 2015; Kelty-Stephen and Dixon, 2014; Teng et al., 2016). Visual inspection of impulse-response plots indicated rare instances of effects from or responses from CoM-*H*_fGn_ on or to CoP-*SD* (1 trial in the 50-, 135-, and 305-cm conditions), but the regression modeling indicated no stable relationship. This absence of significant impulse-response functions for CoM-*H*_fGn_ on CoP-*SD* is puzzling. However, past evidence suggests that fluctuations in the upper body moderate visual information beyond and possibly in collaboration with microsaccades. CoP and movements of the upper extremities exhibit a close mutual predictive relationship in fractal and multifractal fluctuations even without involving a significant role of torso fluctuations (Mangalam et al., 2020a, 2020b). Indeed, should tensegrity-themed metaphors for the movement system be apt (Turvey & Fonseca, 2014), then we can expect relatively less local relationships, and CoM may be one of the multiple intermediary links in the anatomical change that need not always participate in controlling posture. Fuller-body set of measurements may allow clearer portrayal of causal relationships knitting retinal fluctuations with CoP fluctuations.

## 5. Conclusions and future directions

*T*his work exemplifies that visual-postural dynamics raises new questions only definable at the boundaries among physics of complex systems, psychology, and neuroscience (Bronstein, 2019). This work may inform rehabilitation strategies with clinical populations that benefit from visual control of posture (e.g., patients with Parkinson’s disease). For instance, understanding what form of postural fluctuations support stable visually-guided posture could clarify what form of variability should be built into visual information presented by headmounted displays in therapeutic sessions (Lei et al., 2019) that could fit neatly within decades-long discussions on when or how stimulation with monofractal-or-not variability might offer useful clinical applications (Ducharme & van Emmerik, 2018; Kelty-Stephen & Dixon, 2013; Priplata et al., 2002; Priplata et al., 2003; Rhea et al., 2014; Sotirakis et al., 2020; Vaz et al., 2020).

Recent work has examined the role of the auditory stimulus with elaborations beyond fractal structure in supporting gait (Hove et al., 2012; Hunt et al., 2014; Rodger and Craig, 2016; Uchitomi et al., 2013). Despite never explicitly naming ‘multifractal’ structure as its focus, these studies investigating the auditory stimuli propose to build gait-scaffolding stimulation from the foundation of biological patterns (e.g., gait and music) that marry multi-scale aspect of fractal fluctuations with complex phase relationships amongst participating timescales. This blend of features guarantees that these structures show multifractality (Muñoz-Diosdado, 2005; Telesca and Lovallo, 2011). Likewise, if we study the postural system to identify factors that can minimize sway, it becomes evident that the postural system itself exhibits a multifractal structure. Indeed, the stochastic resonance that the scientific community has been trying to build might be more accessible by using multifractal stimulation. The limitation of fractal temporal correlations is that fractal models are all too symmetrical, and postural sway is known to be intermittent (Lee et al., 2018, 2019). Monofractal stimulation may be ill-suited to the multifractal landscape of postural control. Future work could benefit from considering what multifractal configurations of posture will be most receptive to stimulation, and multifractal geometry may offer the plainest framework for tuning stimuli to multifractal control of posture. In pointing to the importance of variation in fractal temporal correlations, this work is a step towards understanding the multisensory aspect of posture and providing multisensory support for its clinical support and rehabilitation.

In the longer theoretical run, we aim to align this multifractal perspective with the ecological notion that the perceiving-acting organism is an integral whole (Gibson, 1966; Stoffregen et al., 2017). We feel that the tensegrity approaches noted above align with this perspective. Hence, our references to intermodality do not rely on a deeper premise of partitioning a global array of stimulation into piecemeal Aristotelian senses. Instead, we envision that multifractal geometry is an apt modeling framework that respects the fundamental continuity of a measured system. In short, multifractality allows us to model the diversity of the movement system as it encounters a multifaceted layout of surfaces, and while our experimental design adds or subtracts a manipulated factor (e.g., visual precision demands or closing eyes), multifractal modeling allows investigating the measured behavior without assuming that the measured behavior is itself the addition or subtraction of causal factors. On the contrary, any seeming ‘subsystems’ within the perception-action system (e.g., eyes and hands for visual and haptic subsystems, respectively) might be sooner better understood through the same sort of nesting that links the whole organism to context. We imagine that different tissues within the body have different forms allowing different sorts of behaviors, but those different tissues contribute meaning to perception-action through their nesting in the organism. Hence, the way that a unitary perception-action system finds through the task environment thrives from nesting relationships within the organism and at larger scales extending beyond the organism (Kelty-Stephen, 2017). Modeling nesting relationships across many different scales at once is exactly what multifractal geometry accomplishes, and so it may be uniquely relevant to the goal of the ecological approaches to perception-action.

## Data accessibility

All data analyzed in the present study are available upon request.

## Author contributions

D.G.K-S., I.-C.L., K.M.N., and M.M. conceived and designed research; I.-C.L. performed experiments; D.G.K-S., N.S.C., and M.M. analyzed data; D.G.K-S. and M.M. interpreted results of experiments; D.G.K-S. and M.M. prepared figures; D.G.K-S. and M.M. drafted manuscript; D.G.K-S. and M.M. edited and revised manuscript; D.G.K-S., I.-C.L., N.S.C., K.M.N., and M.M. approved final version of manuscript.

## Electronic supplementary material

**Dataset S1.** Data used for mixed-effect regression modeling of impulse-response functions relating CoM-*H*_fGn_ CoP-SD, and CoP-*H*_fGn_

**Dataset S2.** Data used for mixed-effect regression modeling of impulse-response functions incorporating CoP-*t*_MF_ along with the prior set of CoM-*H*_fGn_, CoP-SD, and CoP-*H*_fGn_

**Table S1.** Full output for mixed-effect regression modeling of impulse-response functions relating CoM-*H*_fGn_, CoP-SD, and CoP-*H*_fGn_

**Table S2.** Full output of mixed-effect regression modeling of impulse-response functions incorporating CoP-*t*_MF_ along with the prior set of CoM-*H*_fGn_, CoP-SD, and CoP-*H*_fGn_

